# Correction of non-random mutational biases along a linear bacterial chromosome by the mismatch repair endonuclease NucS

**DOI:** 10.1101/2023.12.20.572499

**Authors:** Oyut Dagva, Annabelle Thibessard, Jean-Noël Lorenzi, Victor Labat, Emilie Piotrowski, Nicolas Rouhier, Hannu Myllykallio, Pierre Leblond, Claire Bertrand

## Abstract

The linear chromosome of *Streptomyces* exhibits a highly compartmentalized structure with a conserved central region flanked by variable arms. As double strand break (DSB) repair mechanisms play a crucial role in shaping the genome plasticity of *Streptomyces*, we investigated the role of EndoMS/NucS, a recently characterized endonuclease involved in a non-canonical Mismatch Repair (MMR) mechanism in archaea and actinobacteria, that singularly corrects mismatches by creating a DSB. We showed that *Streptomyces* mutants lacking NucS display a marked colonial phenotype and a drastic increase in spontaneous mutation rate. *In vitro* biochemical assays revealed that NucS cooperates with the replication clamp to efficiently cleave G/T, G/G and T/T mismatched DNA by producing DSBs. These findings are consistent with the transition-shifted mutational spectrum observed in the mutant strains and reveal that NucS-dependent MMR specific task is to eliminate G/T mismatches generated by the DNA polymerase during replication. Interestingly, our data unveil a crescent-shaped distribution of the transition frequency from the replication origin towards the chromosomal ends, shedding light on a possible link between NucS-mediated DSBs and *Streptomyces* genome evolution.

**GRAPHICAL ABSTRACT:** 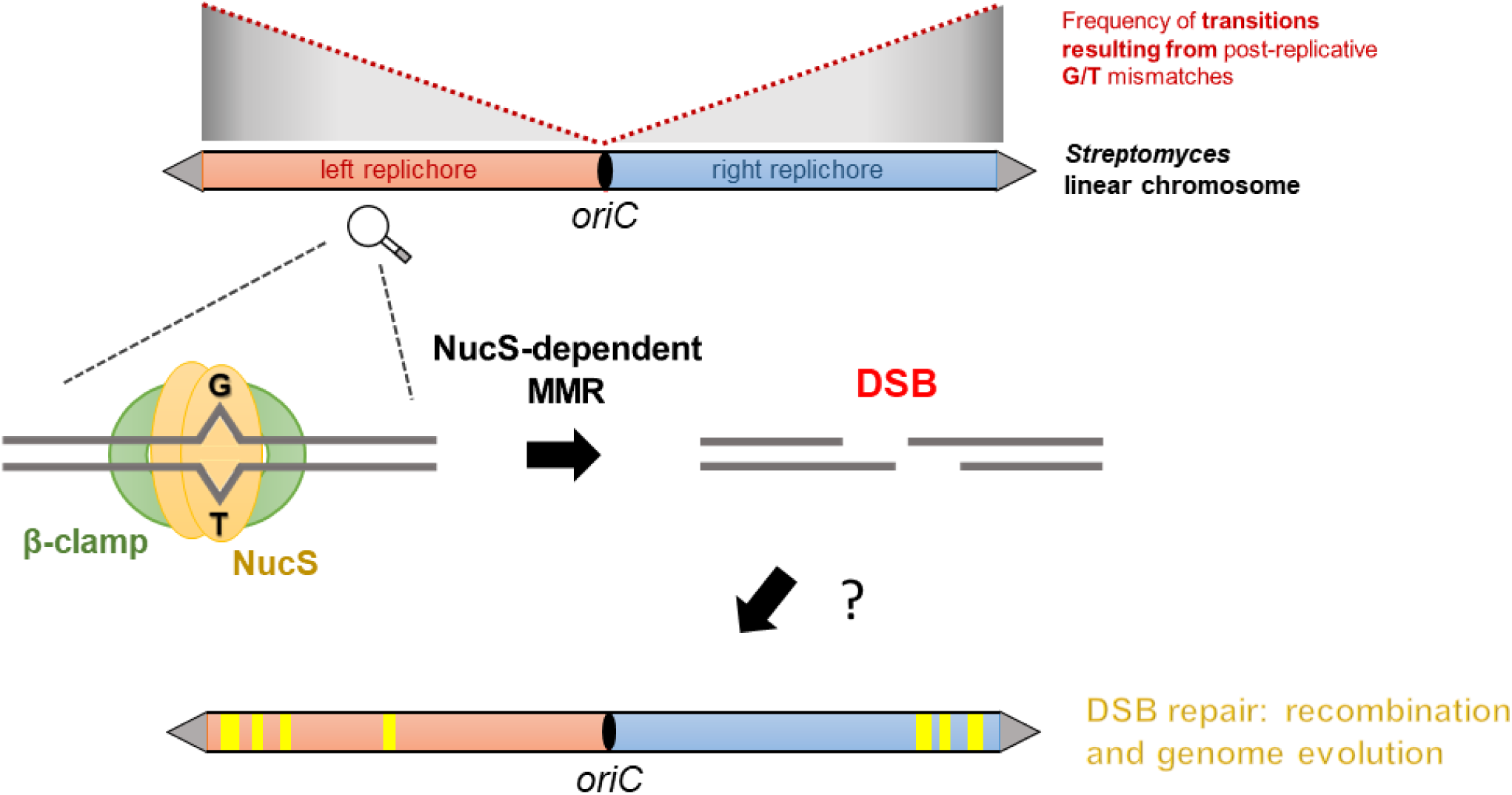

## INTRODUCTION

In all organisms, the fidelity of DNA replication is crucial for the accurate transmission of genetic information. During replication, despite the accuracy of the DNA polymerase, base-pairing errors result in mismatches that are detected and corrected either by the proofreading 3’-5’ exonuclease activity of the DNA polymerase or by the post-replicative MMR (1). The canonical MMR system has been extensively studied in *Escherichia coli* and is well characterized both biochemically and genetically (2). The *E. coli* MMR machinery consists of three major proteins: MutS, MutL and MutH. The process is initiated by MutS which recognizes the mismatch and recruits MutL and MutH proteins. In *E. coli*, the daughter strand is transiently unmethylated during DNA replication, while the parent DNA strand retains its methylation pattern. As a result, the endonuclease MutH nicks the unmethylated strand of the duplex. This provides an entry point for DNA helicases, such as UvrD, to unwind the DNA, allowing many other proteins to take part in the process including single-strand exonucleases which excise a short region of nucleotides from the mismatched strand. The single-stranded DNA gap is then accurately filled-in and sealed by DNA polymerase and DNA ligase, respectively, in order to restore the original genetic information (3). MMR deficient *E. coli* mutants exhibit a 100- to 1, 000-fold increase in mutation rate resulting in a hypermutable phenotype (4). Although the canonical MMR system is widespread, bioinformatic studies surprisingly failed to identify homologs of the *mutSL* genes in the genomes of many archaeal species and of almost all members of the actinobacteria phylum (5). However, these organisms exhibit a mutation rate comparable to that of canonical MMR-bearing bacterial species (6, 7), suggesting the existence of an alternative MMR pathway.

NucS, discovered in the archaeon *Pyrococcus abyssi*, was reported as a novel endonuclease that degrades single-stranded regions on branched DNA (8). Structural analysis of *P. abyssi* NucS revealed a two-domain enzyme that adopts a dimeric conformation. The N-terminal domain is involved in dimerization and DNA binding, while the C-terminal domain carries a minimal RecB-like domain and contains the catalytic site (8). A recent study identified the orthologs of NucS, named EndoMS (mismatch-specific endonuclease) from the hyperthermophilic euryarchaea *Pyrococcus furiosus* and *Thermococcus kodakarensis*, which specifically recognizes double-stranded DNA (dsDNA) containing a mismatch (9). Strikingly, biochemical characterization revealed that EndoMS from *T. kodakarensis* introduces double-strand breaks (DSBs) by cleaving both strands of mismatched substrates. While dsDNA containing G/T, G/G, T/T, T/C and A/G mismatches is cleaved with variable efficiency, no activity on A/C, C/C, and A/A mispairs is detected *in vitro*. This endonuclease activity is considerably enhanced by the association with the sliding clamp PCNA (proliferating cell nuclear antigen). Sliding clamps are found in all organisms and are considered to be a universal platform that recruits numerous DNA-processing enzymes during replication and repair (10, 11). In *E. coli*, the replication clamp interacts with MutS and MutL, as well as with the DNA polymerase III, suggesting a coupling between DNA replication and MMR (12, 13). NucS was in fact historically discovered as an interacting partner of PCNA in *P. abyssi* (14) and this interaction modulates NucS structure and activity (8, 15). *P. abyssi* NucS and *T. kodakarensis* EndoMS exhibit a PCNA-interacting protein (PIP) box in their C-terminal domain (14). A similar box is also known as the clamp-binding motif CBM in bacteria and has a general [EK]-[LY]-[TR]-L-F consensus sequence that tolerates greater variability than the PIP-box (16). The crystal structure elucidation of EndoMS dimer with dsDNA confirmed a preferred binding to specific mismatches and a preliminary complex model with PCNA was proposed (17). This interaction of a restriction enzyme with the sliding clamp is a remarkable specificity that has never been reported before. In actinobacteria, the biological role and the enzymatic activity of EndoMS/NucS (referred to as NucS hereafter) have been mainly studied in *Mycobacterium* species and *Corynebacterium glutamicum*. Deletion of *nucS* in *Mycobacterium smegmatis* and in *C. glutamicum* leads to the increase of spontaneous mutation rates and causes a transition-biased mutation spectrum, highlighting its role in mutation avoidance (18–22). Cleavage assays have shown that NucS of *C. glutamicum*, like archaeal enzymes, generates a DSB by preferentially cleaving G/T mismatched DNA substrate. The activity of the enzyme is dependent on the presence of divalent cation cofactors, Mn^2+^ or Mg^2+^, and its efficiency is greatly enhanced by the addition of β-clamp, the bacterial homolog of PCNA. However, such cleavage has not yet been demonstrated in *Mycobacterium* species. Taken together, these results suggest that NucS corrects mismatches that have escaped the proofreading activity of the polymerase and represents an important component of the non-canonical MMR system. Most importantly, unlike the canonical MMR, NucS creates a DSB that, if left unprocessed, should be dangerous to the cell. DSBs are indeed the most detrimental type of DNA damage in all living cells. They can be repaired by homologous recombination when a DNA template is available or by illegitimate recombination for which a homologous template is not necessary. How the DSB induced by NucS during MMR is repaired and how the correct base pair is restored has yet to be elucidated.

*Streptomyces* are soil bacteria belonging to the phylum of actinobacteria and are known as the most prolific producers of specialized metabolites used in medicine and other industries (23). These bacteria have original genomic features including a high GC content (∼72%) and a large linear chromosome (6-12 Mb) ended by terminal inverted repeats (TIR). The genome of *Streptomyces* is highly compartmentalized with a conserved central region flanked by variable chromosomal arms (24, 25). Preliminary genomic comparison studies revealed that the loss of synteny at the terminal arms results from DNA insertions and deletions (26). Recently, a more comprehensive analysis performed on 125 *Streptomyces* genomes highlighted that the great flexibility of the chromosomal arms is due to gain and loss of genes resulting from horizontal gene transfer or genome shuffling (25). This phenomenon is directly linked to large genomic rearrangements spontaneously occurring in the arms, including large deletions (up to 2 Mb), chromosome circularization and arm replacement events (27–30). Recent work has shown that repair of DSBs in the chromosomal arms resulted in all the chromosomal rearrangements observed in the spontaneous variants (31). DSBs in *Streptomyces* can be repaired by homologous recombination but also by Non-Homologous End-Joining (NHEJ), a widespread illegitimate recombination mechanism among eukaryotes that is not ubiquitous in bacteria (31–34). The repair of DSBs in the central part of the chromosome can be mutagenic when mediated by NHEJ, with deletions of up to 20 kb, whereas repair at the terminal regions can result in huge genomic alterations such as deletions of up to 2.1 Mb, or arm replacement (mediated by a single crossing-over between two homologous sequences present in both arms, or by break-induced replication events). Some scar analyses revealed insertion events of ectopic sequences ranging from tens to thousands of nucleotides, highlighting possibilities of potential integration of foreign DNA in a natural environment. Therefore, DSB repair has been proposed as a driving force for the huge plasticity and evolution of *Streptomyces* genome. Various exogenous sources such as UV light, ionizing radiation or mutagenic chemicals can induce DSBs, but vast amount of these are thought to arise from internal cellular processes, for instance during DNA synthesis when the replication fork collapses. The recent discovery of the non-canonical MMR unveils a new putative internal origin of DSBs that could have a role in *Streptomyces* genome plasticity. In this study, we used mutation accumulation experiment to demonstrate the role of NucS in mutation avoidance in *Streptomyces ambofaciens* and we showed the enzyme cleavage activity on diverse base pair mismatches, particularly on G/Ts, generating a DSB. Our results notably highlight a transition bias that increases from the replication origin towards the extremities of the chromosome of strains lacking NucS, suggesting that the non-canonical MMR system is more active in the terminal regions of *Streptomyces* linear chromosome.

## MATERIAL AND METHODS

### Strains, plasmids, and growth conditions

DH5α *E. coli* strain was used as the cloning host for plasmid construction. Plasmid pGEM®-T Easy (Promega) was used as a primary cloning vector. *E. coli* BL21 (DE3) strain was used for *nucS* and *dnaN* heterologous expression. Plasmid pET15B is the expression vector used in this study, and pSBET is a helper plasmid for heterologous expression. *E. coli* ET12567 pUZ8002 was used as a donor for intergeneric conjugation to introduce DNA constructs into *S. ambofaciens.* Plasmid pUZ8002 carries the conjugative functions. *E. coli* strains were grown in Luria-Bertani (LB) medium at 37°C, except for the BW25113 pKD20 thermosensitive strain used for PCR targeting which was grown at 30°C. Plasmid pKD20 expresses the phage λ Red recombination system. The pOSV508 plasmid allowing the expression of *int* and *xis* from pSAM2 (35) is used in *Streptomyces* to excise genetic cassettes by site-specific recombination. The integrative plasmid pSET152 is used for complementation in *Streptomyces.* The *Streptomyces* mutants mentioned in this study were derived from our reference strain *S. ambofaciens* ATCC23877. *S. ambofaciens* strains were grown on Soya Flour Mannitol agar (SFM) for spore harvest. Single colony isolation for phenotypic observation and growth for DNA extraction were performed on Hickey-Tresner (HT) agar and liquid medium, respectively.

### Deletion of nucS in S. ambofaciens

The *nucS*-deficient mutants were generated by PCR targeting (36). Briefly, the deletion was carried out in a recombinant bacterial artificial chromosome (BAC) containing a 35 kb region containing *nucS* gene, which was replaced by an *oriT-aac*(*3*)*IV* apramycin resistance cassette using the lambda-red system in the highly recombinogenic *E. coli* BW25113/pKD20 strain. This recombinant BAC was then conjugated into *S. ambofaciens* using, as a donor strain, the non-methylating *E. coli* ET12567 which harbors the mobilizing pUZ8002 plasmid. Double-crossover events leading to the replacement of genomic *nucS* by the cassette were monitored by screening for both apramycin resistance and kanamycin sensitivity (selection of the loss of the BAC vector). As the disruption cassette is flanked by pSAM2 *attL* and *attR* sites, its excision by site-specific recombination can be induced thanks to the introduction of pOSV508 plasmid allowing the expression of *int* and *xis* from pSAM2 (35). The excision event was verified by PCR in the exconjugants. The Δ*nucS*-1 and Δ*nucS*-2 mutants originate from two well-isolated colonies of the same conjugation experiment, whereas Δ*nucS*-3 and Δ*nucS*-4 mutants originate from a second conjugation. These two conjugations were performed with two modified BACs. The resulting four *nucS*-deficient strains were therefore considered as independent.

### Complementation of Δ*nucS* mutants

*S. ambofaciens nucS* locus (SAM23877_5135) including the promoter and the coding region was amplified by PCR using primers nucS_compl_F/R containing an EcoRI restriction site (Supplementary Table S1). The PCR fragment was ligated into the pGEM-T Easy vector, sequenced and then released by EcoRI digestion in order to be inserted into the conjugative and integrative plasmid pSET152, resulting in the pSET152-*nucS* complementation vector. The latter was introduced into the four Δ*nucS* mutants by intergeneric conjugation as described above. As pSET152 plasmid is integrative at the φC31 chromosomal *attB* site, there is only one complementation copy of *nucS* gene per chromosome. The integration of the pSET152-*nucS* plasmid in the chromosome was verified by PCR.

### Estimation of mutation rate by fluctuation tests

Spontaneous mutation rates were estimated using fluctuation assays (37). Isolated colonies of wild-type (WT) and Δ*nucS*-2 strains were excised after 7 and 5 days of growth on SFM at 30°C, respectively, and vortexed vigorously in water to homogenize spores in solution. Three volumes of 100 µL of each suspension were plated on three SFM plates supplemented with rifampicin (50 µg/mL) and appropriate dilutions were plated on SFM in order to determine the total viable cell number. Rifampicin-resistant colonies were counted after 4 days of incubation at 30°C. The mutation rate was determined using the Ma-Sandri-Sarkar maximum likelihood method (38) and confidence intervals were calculated according to the literature (39). These calculations were performed using the FALCOR web tool available at https://lianglab.brocku.ca/FALCOR/ (40).

### Heterologous expression and purification of His-tagged NucS*_Sam_* and β-clamp_Sam_ proteins

The coding sequences of *nucS* (SAM23877_5135) and of *dnaN* (SAM23877_3780), which code for NucS_Sam_ and β-clamp_Sam_, respectively, were amplified by PCR from *S. ambofaciens* chromosome. The amplification of *nucS* and *dnaN* was carried out with the primer couples nucS_F/R and dnaN_F/R, respectively (Supplementary Table S1). The PCR fragments were ligated into the pGEM®-T Easy vector and were verified by sequencing. The digestion by NdeI and BamHI released *nucS* and *dnaN* sequences, which were further cloned into the pET15b expression vector. Using these constructs, a His-tag is added to the N-terminal end of NucS_Sam_ and β-clamp_Sam_. The *E. coli* BL21 (DE3) strain, containing the helper plasmid pSBET (41), was used for protein expression. Expression and purification of NucS_Sam_ and β-clamp_Sam_ were performed using the same procedure. Bacteria were grown at 37°C to exponential phase in LB medium supplemented with 50 µg/mL kanamycin and 100 µg/mL ampicillin. The expression of chaperones was boosted by adding ethanol to the bacterial culture to a final concentration of 0.5% and by cooling it at 4°C for 3 h. IPTG was then added to a final concentration of 100 µM and cells were further grown for 20 h at 20°C. Cells were collected, diluted in buffer A (30 mM pH 8.0 Tris-HCl, 200 mM NaCl, 10 mM imidazole, 0.05% polyethyleneimine) and disrupted by sonication. The soluble extract was applied to an IMAC-Select affinity gel column (Sigma Aldrich, St. Louis, MO, USA) equilibrated in buffer A. After extensive washing, the protein was eluted using the same buffer A with an imidazole concentration of 100 mM. Fractions containing the protein of interest were pooled, concentrated using an Amicon centrifugal filter (Millipore) with a 10 kDa cut-off and loaded onto a Superdex 200 pg HiLoad 16/600 (GE Healthcare, 1 mL/min). Sample purity was assessed by 12% SDS-PAGE analysis and sample concentration was determined by spectrophotometry using a theoretical molar extinction coefficient at 280 nm of 20, 190 M^−1^.cm^−1^ and 19, 940 M^−1^.cm^−1^ for NucS_Sam_ and β-clamp_Sam_, respectively.

### Cleavage assay

Heteroduplex DNA was prepared by annealing oligonucleotides at 95°C for 10 min. The sequence and the different combinations of oligonucleotides are shown in Supplementary Table S2. The cleavage reaction of the dsDNA substrate was generally performed at 30°C for 60 min in a buffer containing 2.4 µM NucS_Sam_ and/or 1.2 µM β-clamp_Sam_, 25 mM Bis-Tris pH 6.4, 2.5 mM MnCl_2_, 0.1% Triton X-100. For cleavage reaction performed with Mg^2+^, either 2.5 mM MnCl_2_ was replaced by 2.5 mM MgCl_2_ in the above conditions, or the reaction conditions described by Takemoto *et al.* (21) were used as follows, 20 mM Tris–HCl, pH 8.0, 2.5 or 10 mM MgCl_2_, 4 mM dithiothreitol, 0.071 mg/ml BSA, and 30 or 100 mM K[oAc]. Monomeric NucS_Sam_ fraction was used in these assays. The products were analyzed by 10% native PAGE. At least three biological replicates were done for each mismatch configuration.

### Mutation accumulation experiments

The WT strain and the four independent *nucS*-deficient mutants were used for mutation accumulation (MA) experiments. Three lineages per Δ*nucS* mutants and twelve WT lines were generated. Each line was generated by streaking a single colony. A sporulation cycle was initiated by streaking a single well-isolated colony from each lineage onto a new SFM plate and incubating at 30°C for 7 days. This procedure was repeated every week for 60 cycles. To keep track of evolved lineages, colonies streaked at each new cycle were picked from the agar plate, resuspended in 20% glycerol and stored at -20°C.

### Genomic DNA preparation and whole genome sequencing analysis

Parental and evolved strains at the end of the 60^th^ sporulation cycle were grown in liquid HT medium to extract genomic DNA (42). Whole genome sequencing was performed using an Illumina Genome Analyzer to obtain 300-bp paired-end reads (Mutualized Platform for Microbiology (P2M), Pasteur Institute, Paris). Data analysis was performed using Geneious Prime 2023.2.1. Reads from each lineage were trimmed based on their quality (Minimum quality = 20; primers, indexes and reads less than 20 bp in length were removed). Trimmed reads from parental strains only were mapped to the *S. ambofaciens* ATCC23877 reference genome (GenBank: CP012382.1) using Geneious mapper with the following parameters: sensitivity = “medium-low”, fine tuning = “iterate up to 5 times”, minimum mapping quality = “30”, only map paired reads which = “map closely nearby”. The mean depth coverage of each assembly ranged from 113 to 481x. Consensus sequence was generated from alignments using the highest quality parameter as a threshold and “N” was assigned to sites with coverage < 20. Annotation was performed with Prokka (43) using a custom database containing all the genes from *S. ambofaciens* ATCC 23877 chromosome. These annotated genomes were then used as references for mapping trimmed reads from evolved lines. The same Geneious alignment settings as above were applied. The mean depth coverage of each mapping ranged from 45 to 565x. Variant calling was performed using the in-built “Find Variations/SNPs” feature. A single-nucleotide polymorphism (SNP) was called if the position was covered by at least 20 reads, was found at a frequency of at least 0.9, and was not found in a niche (>3 SNPs in a sliding 100-bp window with a 1-bp step size). Note that all the genomic analyses were done on the whole genome of *S. ambofaciens* ATCC23877 with the exception of the 89 kb circular plasmid pSAM1 and repeated sequences (including the 200 kb long TIR of *S. ambofaciens*). Therefore, in the manuscript, when we refer to the chromosome, it is without the above sequences.

### Genomic comparison of *Streptomyces* close environmental strains

Genomes of RLB1-8 and RLB3-17 (accession numbers CP041650 and CP041610, respectively) environmental strains (44) were used in this study. ANIb (Average Nucleotide Identity using BLAST) was calculated between both genomes: we averaged the two reciprocally obtained ANIb scores (45). For transition analysis in whole genome, both sequences were aligned using the progressive Mauve algorithm (46). The SNP call table was exported with MAUVE tool ‘export SNPs’. For transition analyses in coding sequences, both genome sequences were annotated using RAST (47), with the RAST pipeline (https://rast.nmpdr.org/) using RASTtk (v1.3.0) with all parameters by default. After extraction, RAST annotations (with GenBank files) were formatted to a usable format for the subsequent pipeline. BLASTp was performed between each gene coding for a protein of one genome against that of the other genome and reciprocally. Orthologs were defined by identifying BLASTp reciprocal best hits (BBH) with at least 40% identity, 70% coverage, and an E-value lower than 1e-10. For each orthologous gene pair, the nucleotide sequences of both genes were retrieved and aligned with MAFFT (48). In these alignments (one per orthologous gene pair), identified non-gap differences corresponded to SNPs. RLB1-8 genome was used as reference. The sequence of the 5 identified integrative and conjugative elements present in RLB1-8 genome (49) were removed from the transition analysis to avoid bias caused by their possible redundant nucleotide sequences. The 357 kb TIR of RLB1-8 genome was also excluded, as in the MA study.

### Statistical analyses of mutation distribution along the chromosome

For the analyses of mutation distribution along the chromosome, the number of mutations was counted within a sliding non-overlapping 100 kb window, starting from the replication origin *oriC*, extending towards the left or the right end of the chromosome. The number in windows of the right replichore was added to the number in windows at the same distance of *oriC* in the left replichore. The sum was represented as a function of the distance from *oriC.* As the left replichore is slightly longer than the right one in both *S. ambofaciens* ATCC23877 and RLB1-8 genomes, values corresponding to the last 100 kb windows at the extremity of the left replichore were excluded from the analyses. Correlation analyses between the number of mutations and the distance from the origin of replication in the genomes of evolved lines or between the genomes of environmental strains were done with R software (https://www.R-project.org/). Pearson correlation test was used when the distribution of the residues was compatible with the normal low and Kendall’T coefficient was calculated when the distribution of the residues was not compatible with the normal low.

### Reagents

All cloning reagents, including restriction enzymes (FastDigest), T4 DNA ligase, and DNA polymerases (DreamTaq, Phusion) were provided by Thermo Fisher Scientific. All molecular biology kits (PCR purification, gel extraction and plasmid miniprep) were also provided by Thermo Fisher Scientific.

## RESULTS

### NucS is highly conserved in *Streptomyces*

The persistence of *nucS* across *Streptomyces* genus was determined by searching for *nucS* orthologs in the genomes of a collection of 125 species representative of the genus diversity (25). A single copy of *nucS* was present in all species and, as every core gene, is localized in the central region of the chromosome. NucS protein sequence consists of 223 amino acids and is highly conserved with identities ranging from 86% to 97% when using NucS from *S. ambofaciens* (NucS*_Sam_*) as a reference. The ubiquity of *nucS* through *Streptomyces* genus suggests its important role in cell development and survival in *Streptomyces*. When compared to NucS from *M. smegmatis* and *Mycobacterium tuberculosis* (18), or from *C. glutamicum* (20) the identity drops to 75%, 78%, and 72% respectively, highlighting the significant conservation of the protein among actinobacteria. NucS*_Sam_* shares structural homologies with NucS from *P. abyssi* (8), bearing an N-terminal DNA binding domain connected by a linker to the C-terminal RecB-like nuclease domain. The well-known RecB motifs for nuclease activity are present in NucS*_Sam_* and most importantly all the catalytic residues are conserved across actinobacteria (Supplementary Figure S1). The consensus clamp-binding KLRLF motif is also present in the C-terminal part of all *Streptomyces* NucS proteins. These data strongly suggest that NucS*_Sam_* may have a similar activity to NucS*_Pab_*.

### NucS*_Sam_* is involved in mutation avoidance

To assess the effect of *nucS* deletion, four independent mutants deleted for *nucS* (Δ*nucS*) were generated by PCR targeting method in *S. ambofaciens* ATCC 23877 strain. For each independent Δ*nucS* mutant, a complemented strain, expressing *nucS* WT copy under the control of its own promoter, was obtained using the integrative plasmid pSET152. The Δ*nucS* mutants were grown for 8 days at 30°C on HT solid medium for phenotype analysis. Compared to the WT, all *nucS*-deleted strains shared the same remarkable phenotype consisting of a high frequency of sectored colonies in their progeny (Figure 1A and B). This phenotype is heritable. The proportion of sector-harboring colonies was quantified in the 4 independent Δ*nucS* mutants, in the WT and in the complemented strains. The occurrence of sectored colonies was significantly greater in Δ*nucS* strains compared to the WT strain (overall 82% versus 7% respectively; Z-test for proportions with Bonferroni correction, P < 10^−3^; Figure 1C). In the complemented strains, this frequency was drastically reduced to levels comparable to those of the WT (on average 9% *vs* 7% respectively). This indicates that expressing *nucS* behind its original promoter restored the WT phenotype. In the WT background, it has been shown that the spontaneous variability of the colonial phenotype originates from mutations, affecting phenotypical traits in *Streptomyces* such as pigmentation or sporulation (27, 50). We therefore assumed that the specific phenotype observed for *nucS*-defective strains was the result of mutation accumulation.

**Figure 1.**
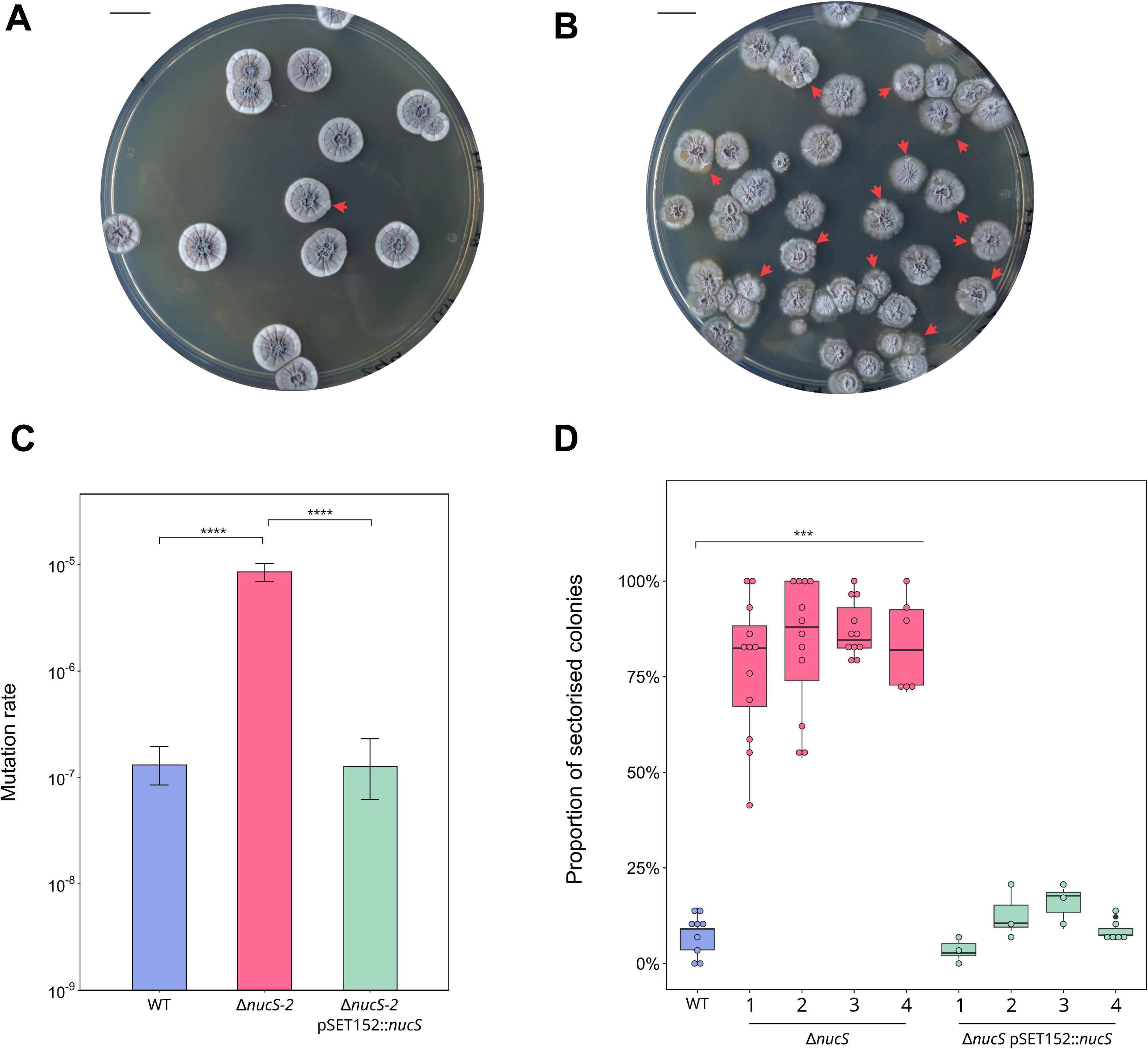
Colonial phenotype of *nucS-*deleted mutants. (A) *S. ambofaciens* WT strain and (B) Δ*nucS* mutant were grown on HT agar for 8 days at 30°C. A sectorized colonial phenotype was observed for *nucS-*deleted strain (red arrows). Scale bar represents 1 cm. (C) Proportion of sector-harboring colonies counted in the WT, the 4 independent Δ*nucS* mutants and in their complemented strains denoted as Δ*nucS* pSET152::*nucS* (z-test for proportions with Bonferroni correction, ***P<0.001). (D) The rate of spontaneous mutations conferring rifampicin resistance was determined by fluctuation assay in WT, Δ*nucS*-2 and its complemented strain Δ*nucS-2* pSET152::*nucS*. Error bars represent 95% confidence intervals.

To confirm that *nucS* deletion results in a hypermutator phenotype, the spontaneous emergence of rifampicin-resistant colonies was monitored by fluctuation assays. Originally described by Luria and Delbruck (37), fluctuation tests are widely used for estimating mutation rates. As all 4 Δ*nucS* strains exhibited the same colonial phenotype, Δ*nucS*-2 was selected for further assays. The same procedure was applied to the corresponding complemented strain. Deletion of *nucS* resulted in an increase in mutation rate by a 130-factor compared to the WT strain (1.63 × 10^−6^ *versus* 1.24 × 10^−8^ mutations per cell, respectively, Figure 1D). Of note, a mutation rate comparable to that of the WT strain was recovered in the complemented strain (1.24 × 10^−8^ *versus* 1.08 × 10^−8^ mutations per cell, respectively) confirming that the absence of NucS was responsible for the observed increase in the mutation rate. Similar findings were reported in *Streptomyces coelicolor*, in which inactivation of *nucS* resulted in a 2-log increase in mutation rate, although no colonial phenotype was reported in this study (18). Altogether, these results showed that *nucS* inactivation caused a hypermutator phenotype and highlighted the key role of NucS in mutation avoidance.

### Purification of His-tagged NucS*_Sam_* and His-tagged β-clamp*_Sam_* proteins

NucS from the actinobacterium *C. glutamicum* has been recently demonstrated as capable of cleaving mismatched dsDNA (20, 21). Yet, Castañeda-García and colleagues (18) have shown that NucS from *M. smegmatis* has ssDNA-binding activity but no cleavage activity was detected neither on ssDNA nor on mismatched dsDNA. For this reason, the activity of NucS*_Sam_ in vitro* was investigated. To achieve this, we purified N-terminal His-tagged NucS*_Sam_* by cloning the *nucS* coding sequence (SAM23877_5135) in the pET15b expression vector for heterologous expression in *E. coli* BL21. The fusion protein was purified using affinity chromatography and gel filtration. Based on the amino acid sequence, the calculated molecular weight of the protein is 24, 762.44 Da. The gel filtration chromatogram showed a main peak at an elution volume corresponding to an estimated molecular weight of 41 kDa (Supplementary Figure S2). Additional peaks were observed with estimated molecular weights of 61 and 115 kDa, indicating the presence of multimeric forms. Because it was difficult to firmly conclude about the oligomerization state of the major form of purified NucS*_Sam_*, the fractions corresponding to the main peak were collected and subjected to size exclusion chromatography coupled with multi-angle light scattering (SEC-MALS). The analysis revealed two main peaks with molecular weights of 30, 026 Da and 61, 151 Da, which would correspond to monomeric and dimeric forms of NucS*_Sam_*, respectively. Because the homogeneity of these peaks was moderate, molecular weights were determined considering the left side of the peaks. The gel filtration profile and SEC-MALS analysis indicate that the protein may exist in an equilibrium between a monomeric and dimeric state, with the monomeric form being more prevalent under the purification conditions employed in this work. This observation may also suggest that a DNA substrate is necessary for monomers to assemble into a dimer. The processivity-promoting factor β-clamp from *S. ambofaciens* (β-clamp*_Sam_*), which is encoded by the *dnaN* gene (SAM23877_3780), was also purified following the same protocol as the one used for NucS*_Sam_*. Its calculated molecular weight is 39, 882.35 Da. The gel filtration chromatogram revealed a single peak with an estimated molecular weight of approximately 97 kDa while SEC-MALS indicated a molecular weight of approximately 84 kDa. This highlighted the homodimeric nature of β-clamp*_Sam_* in solution and confirmed previous results obtained for β-clamp from *C. glutamicum* and *T. kodakarensis* (9, 20).

### NucS_Sam_ mismatch-specific endonuclease activity results in DSB formation *in vitro*

Cleavage assays were performed *in vitro* to test whether purified NucS*_Sam_* possesses an endonuclease activity on mismatched dsDNA. The DNA substrate used in this work was a 43 bp-long dsDNA containing different combinations of mismatched base pairs at the 21^st^ nucleotide pair. One strand of the substrate was fluorescently labeled at the 5’ end with 6FAM dye. As a control, a dsDNA substrate with no mismatch was also subjected to NucS*_Sam_* activity. The cleavage products generated by NucS*_Sam_* were analyzed qualitatively using a 10% native PAGE. No cleavage product was observed when using the control substrate. NucS*_Sam_* displayed cleavage activity on substrates containing G/T, T/T, and G/G mismatches. The cleaved product co-migrated with a control dsDNA containing a 5’ protrusion of a five nucleotide-single-stranded DNA and expected to correspond to the product after a cleavage within the mismatch (data not shown). No activity was detected with substrates containing T/C, A/A, A/C, G/A, or C/C mismatches (Figure 2A shows the results of one of the three biological replicates). Notably, the addition of β-clamp*_Sam_* to the reaction greatly enhanced the cleavage efficiency of NucS*_Sam_*, as evidenced by the presence of a more intense band corresponding to the cleaved substrate. The presence of β-clamp*_Sam_* did not alter the substrate specificity of NucS*_Sam_*. The presence of the metallic enzymatic cofactor Mn^2+^ was always required for NucS*_Sam_* cleavage activity but no activity was observed with Mg^2+^ cofactor (data not shown).

**Figure 2.**
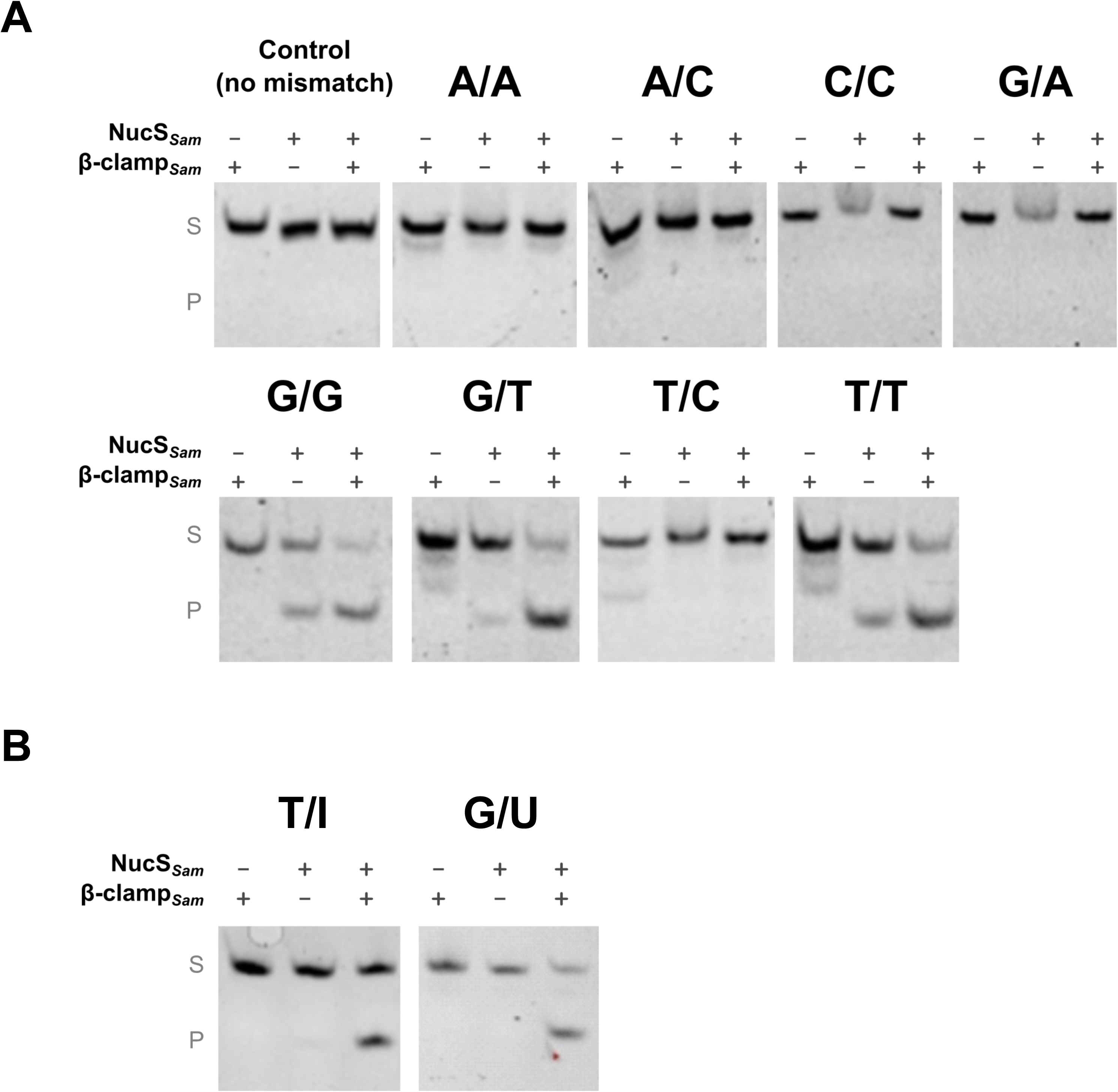
NucS*Sam* mismatch specific cleavage activity. (A) *In vitro* cleavage assays were performed with 6-FAM-labeled dsDNA substrates (43-bp, 50 nM) containing no mismatch (control) or single base-pair mismatches (A/A, A/C, C/C, G/A, G/G, G/T, T/C or T/T) or (B) single deaminated bases (I instead of A or U instead of C). Each substrate was incubated for 60 min at 30°C, either with 2.4 µM of NucS*Sam* or with 1.2 µM of β-clamp*Sam* or with both proteins at the above concentrations. Products were separated by 10% native PAGE. Substrates and cleavage products are indicated by “S” and “P” letters on the left side of the panels. The figure is representative of three experiments.

Recently, an interesting study (51) showed that NucS from the euryarchaeon *Thermococcus gammatolerans* is able to cleave dsDNA containing deaminated bases such as uracil and hypoxanthine. Deamination process consists in the removal of an amino group from a molecule. In cells, deamination of DNA bases can occur spontaneously and produce mutagenic DNA lesions. Specifically, the removal of the amino groups from cytosine and adenine results in the formation of uracil (U) and hypoxanthine (I), respectively. If not repaired, these lesions can cause transition mutations since uracil pairs with adenine and hypoxanthine pairs with cytosine. We assessed whether NucS*_Sam_* had an endonuclease activity on dsDNA containing deaminated bases. *In vitro* cleavage tests showed that NucS*_Sam_* alone had no activity against G/U and T/I substrates, in contrast to the results reported by Zhang and collaborators (51) (Figure 2B). Interestingly, the activation of such mismatch-specific endonuclease activity was observed when β-clamp*_Sam_* was added to the reaction. These results are original as they suggest that NucS*_Sam_* participates in a process of DNA repair other than mismatch repair such as the base excision repair pathway.

### Mutation rate is affected in *nucS*-deleted strains

To investigate the *in vivo* activity and specificity of NucS*_Sam_*, a mutation accumulation (MA) assay was conducted. Twelve independent WT lines and three independent lines for each *ΔnucS* mutant were tracked for 60 sporulation cycles. Weekly, a single colony from each lineage was streaked onto fresh growth medium. By performing whole-genome sequencing on both parental and evolved strains, we identified the mutations that occurred throughout the MA experiment. To ensure accurate mapping, a rigorous approach was implemented by excluding repetitive sequences from our analysis (including the 200 kb long TIR of *S. ambofaciens*). As a result, the sequence coverage for each line accounted for more than 89% of the complete *S. ambofaciens* ATCC 23877 genome. A total of 156 mutations were found in the WT lines, corresponding to an overall mutation rate of 2.84 × 10^−8^ mutation per nucleotide per cycle (Table 1). The number of generations occurring during a sporulation cycle of 7 days in *Streptomyces* was estimated to be approximately 24. Thus, the WT mutation rate would be around 1.18 × 10^−9^ per nucleotide per generation, which is comparable to that observed in *M. smegmatis* or *C. glutamicum* (19, 21). The ratio of non-synonymous to synonymous mutations (dN/dS) determined by the MA assay in WT lines was 2.96 (71/24), which was not significantly different from that expected (2.54), based on the codon usage in *S. ambofaciens* ATCC 23877 strain (chi-squared test χ^2^ = 0.4182., *P* = 0.51), suggesting that the selective pressure in our MA protocol was minimal. Overall, the 12 *ΔnucS* lines exhibited a total of 5, 067 mutations (Table 1). The 3 evolved lines derived from each *ΔnucS* independent mutant exhibited an accumulation of mutations ranging from 1, 150 to 1, 372 (Table 1). The deletion of *nucS* resulted in a significant 32-fold average increase in the mutation rate per nucleotide per cycle, with rates of 2.8 × 10^−8^ for the WT and an average of 9.2 × 10^−7^ for the four *ΔnucS* mutants. The ratio of dN/dS mutations in *ΔnucS* lines was 3.13 (3280/1048), which was significantly different from that expected of 2.54 (chi-squared test *P*<10^-5^) but not significantly different from the dN/dS obtained in WT lines (chi-squared test *P*=0.11). Thus, the results obtained with the mutants appear to be close to, but slightly above, the expected result for an unbiased distribution of non-synonymous and synonymous mutations.

**Table 1.** Total number of mutations and mutation rates determined by MA assays.

| Strain | Number of lineages | Average sequence coverage<sup>†</sup> (%) | Total number of mutations | Mutation rate per nucleotide per cycle (x10<sup>-8</sup>) |
| --- | --- | --- | --- | --- |
| **WT** | 12 | 90.23 ±2.49 | 156 | 2.84 |
| ***ΔnucS*** | 12 | 91.34 ±1.78 | 5067 | 92.23 |
| *ΔnucS-1* | 3 | 91.34 ±0.28 | 1372 | 100.64 |
| *ΔnucS-2* | 3 | 92.20 ±0.51 | 1226 | 88.29 |
| *ΔnucS-3* | 3 | 92.70 ±0.41 | 1319 | 94.52 |
| *ΔnucS-4* | 3 | 89.14 ±2.30 | 1150 | 85.39 |

<sup>†</sup> The sequence coverage corresponds to the percentage of the reference genome of *S. ambifaciens* (ATCC 23877: 8,303,940 bp) analyzed for mutation call between evolved lines and their parental strain.

### Mutational spectrum is shifted upon NucS*_Sam_* loss

The mutational profiles and rates of base pair substitutions (BPSs) in *S. ambofaciens* WT and *ΔnucS* lines are shown in Table 2. In the WT MA lines, a total of 147 BPSs was identified, corresponding to a rate of 2.67 × 10^−8^ BPS per nucleotide per cycle. This rate is consistent with the previously reported BPS rate in *S. coelicolor*, which was about 1.5 BPSs per 10^8^ nucleotides per cycle (52). Furthermore, in the 12 *ΔnucS* lines, a total of 5, 048 BPSs was detected, resulting in a BPS mutation rate of 91.88 × 10^−8^ per nucleotide per cycle. This substantial increase represents a remarkable 34-fold difference compared to the WT MA lines. Additionally, our analysis unveiled the presence of 9 indels in the WT lines, ranging from 1 to 4 bp in length, while we detected 19 indels (of 1 to 2 bp) in the absence of NucS*_Sam_*. Therefore, the loss of NucS*_Sam_* did not considerably affect the rate of indels (0.05 × 10^−8^ *vs* 0.16 × 10^−8^ for WT and *ΔnucS*, respectively).

**Table 2.** Spectrum of Base Pair Substitutions (BPSs) and indels.

|  | WT |  |  | *ΔnucS* |  |  |
| --- | --- | --- | --- | --- | --- | --- |
| | Number | Percentage (%) | Mutation rate<br>per nucleotide<br>per cycle ($\times 10^{-8}$) | Number | Percentage (%) | Mutation rate<br>per nucleotide<br>per cycle ($\times 10^{-8}$) |
| **BPSs** | 147 | 100 | 2.67 | 5048 | 100 | 91.88 |
| Transitions | 73 | 49.66 | 1.33 | 4908 | 97.23 | 89.33 |
| A:T>G:C | 32 | 21.77 | 0.58 | 2986 | 59.15 | 54.35 |
| G:C>A:T | 41 | 27.89 | 0.75 | 1922 | 38.07 | 34.98 |
| Transversions | 74 | 50.34 | 1.35 | 140 | 2.77 | 2.55 |
| A:T>T:A | 5 | 3.55 | 0.09 | 12 | 0.24 | 0.22 |
| A:T>C:G | 6 | 3.55 | 0.11 | 20 | 0.40 | 0.36 |
| G:C>T:A | 18 | 11.35 | 0.33 | 34 | 0.67 | 0.62 |
| G:C>C:G | 45 | 31.21 | 0.82 | 74 | 1.47 | 1.35 |
| **Indels** | 9 | 100 | 0.16 | 19 | 100 | 0.35 |
| Insertions | 3 | 33.33 | 0.05 | 9 | 47.37 | 0.16 |
| Deletions | 6 | 66.67 | 0.11 | 10 | 52.63 | 0.18 |

Regarding the BPSs in the WT, the transition/transversion ratio was 1.01 (49.7% vs. 50, 3% respectively). The most prevalent mutation type was G:C>C:G, accounting for 31.2% of total BPSs. Of note, G:C>A:T and A:T>G:C transitions were the second and third most frequent mutation types (27.9% and 21.8% of the BPSs, respectively). The inactivation of NucS*_Sam_* caused a significant bias in the spectrum towards transitions, which accounted for 97.2% of the substitutions whereas only 2.8% of the BPSs were transversions. Consequently, this resulted in a high transition/transversion ratio of 36.1, which was 36-fold higher than that of the WT lines. In addition, *ΔnucS* lines exhibited a remarkable 67-fold increase in the transition mutation rate, while the transversion mutation rate underwent a modest 2-fold increase. Overall, the rates of mutation for all substitutions were higher in *nucS*-deleted lines compared to WT (Figure 3), but it is noteworthy that the mutation rate of the A:T>G:C and G:C>A:T substitutions increased strikingly in the mutant with 94-fold and 47-fold increases, respectively. Altogether, the deletion of *nucS* resulted in a significant shift in the BPS bias, particularly towards A:T>G:C transitions, which accounted to 59.2% of all observed BPSs.

**Figure 3.**
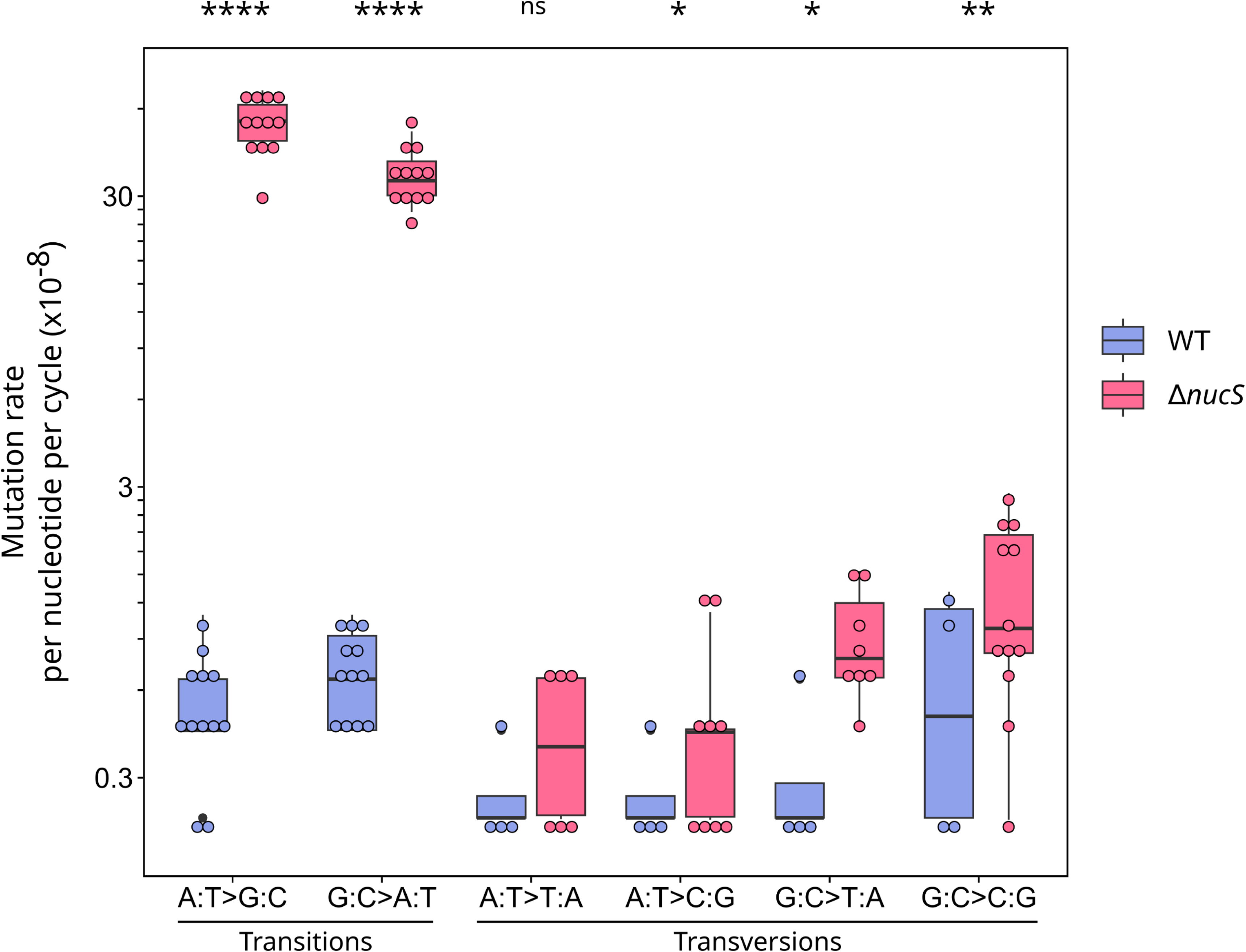
Transition and transversion rates in WT and Δ*nucS* lines. The top of each boxplot indicates 75% percentile, the bottom 25% percentile, and the thick bar inside the box is the median. The minimum and maximum values are shown by the whiskers. Individual data values for each line are represented by circles. Statistical significance levels are denoted as follows *P < 0.05, **P < 0.01, ****P < 0.0001 and ns stands for non-significant (Wilcoxon rank sum test).

### Transition frequency increases towards the extremities of the chromosome

The *S. ambofaciens* chromosome replicates from a centrally located origin of replication towards its ends, defining two replichores of equivalent size. To assess the distribution of transitions along the *S. ambofaciens* chromosome in the twelve evolved lines, the number of transitions was counted within a sliding non-overlapping 100 kb window, starting from *dnaA* gene (at position 4, 021, 374, approximately corresponding to the replication origin *oriC*), extending towards the left or the right end of the chromosome and plotted as a function of the distance from *dnaA* (Figure 4A). A significant positive correlation was observed between the number of transitions and the genomic position for *ΔnucS* lines (Pearson’s correlation coefficient *r=*0.457, *P*=0.004). This suggests that in the absence of NucS, transitions have a crescent-shaped distribution with a minimum at the replication origin of the chromosome. There was no significant correlation for the transition distribution in the chromosomes of the twelve WT lines, probably because the sample size is too small (73 transitions in total). Since 97.3% of all the BPSs are transitions in *ΔnucS* lines, an increase in the BPS number towards the extremities was also observed (Supplementary Figure S3A). When only considering the transitions in coding regions (Figure 4B), the same increasing trend is again noticed (Pearson’s correlation coefficient *r*=0, 468, *P*=0, 003). As a low occurrence of mutations in the central part of the chromosome could be due to negative selection of mutations in essential genes, non-synonymous and synonymous mutations within coding sequences were counted along the chromosome of *ΔnucS* lines and plotted as above. Non-synonymous mutation frequency (Supplementary Figure S3B) increased significantly towards the arm ends (Pearson’s correlation coefficient *r=*0.499, *P*=0.001), but no trend was observed for synonymous mutations (Supplementary Figure S3C). However, the positive correlation found between the non-synonymous mutation number and the genomic position was lost when the non-synonymous mutation number was plotted against the number of synonymous mutations for each position (Supplementary Figure S3D). Indeed, the ratio between non-synonymous and synonymous mutations is homogeneous along the genome, suggesting that there is no clear selection difference depending on the genomic position.

**Figure 4.**
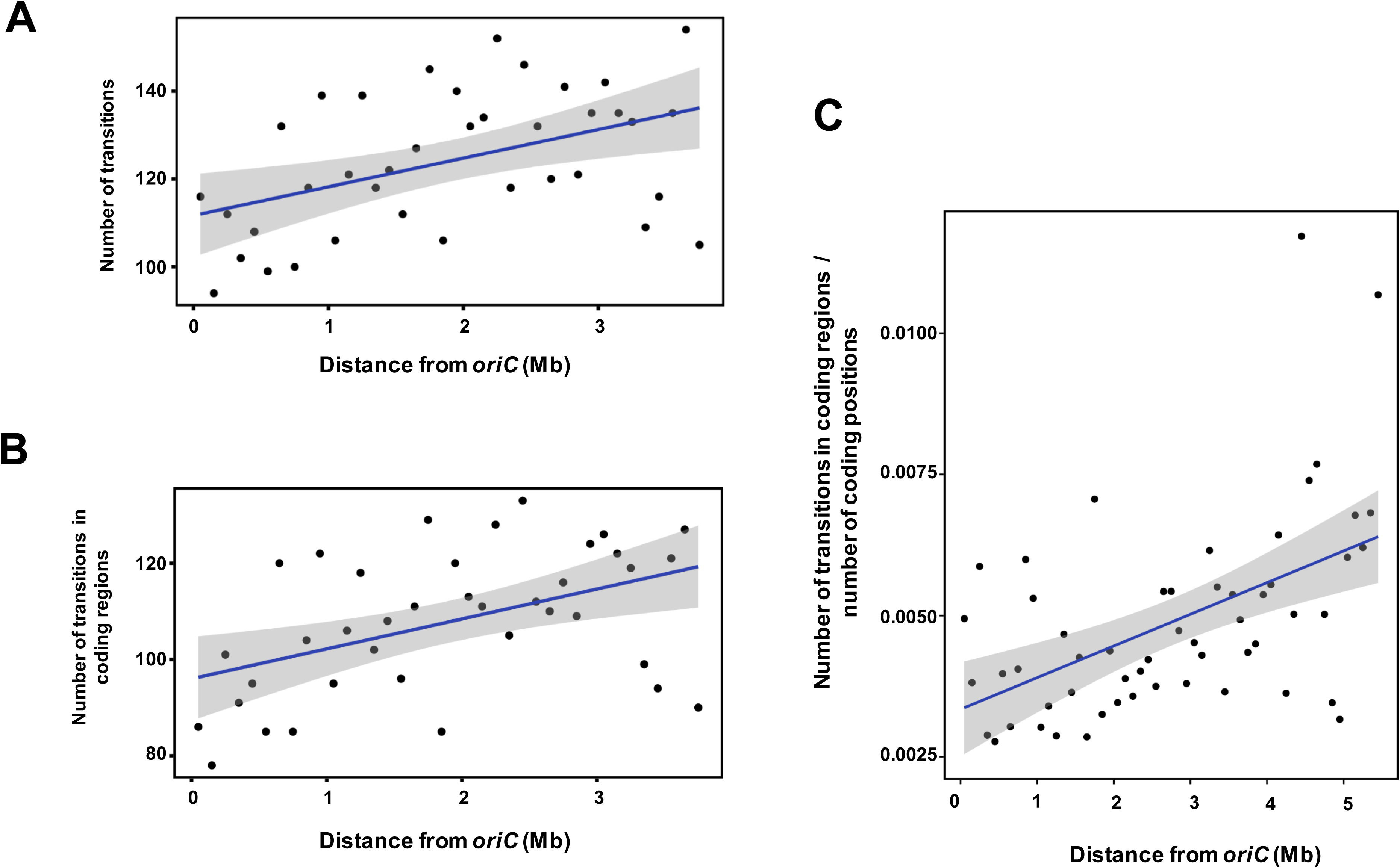
Transition distribution along *Streptomyces* chromosome. (A and B) Transitions along the chromosome of the 12 *S. ambofaciens* evolved Δ*nucS* lines were counted within a non-overlapping 100 kb window, starting from *dnaA* gene (located in the middle of the chromosome at position 4, 021, 374 and approximately corresponding to the origin of replication, *oriC*) and sliding towards the chromosome extremities. The number of transitions in windows of the right replichore was added to the number of transitions of windows at the same distance of *dnaA* in the left replichore. The sum of total transitions (A) or the sum of transitions restricted to the coding sequences (B) was represented as a function of the distance from *dnaA.* Significant positive correlations were observed between the genomic position and the transition count (Pearson’s correlation coefficient *r=*0.457, *P*=0.004) or the number of transitions within the coding sequences (Pearson’s correlation coefficient *r*=0, 468, *P*=0, 003). (C) Transition distribution was estimated along the chromosome of environmental strain RLB1-8 compared to the environmental strain RLB3-17. As above, transitions within coding regions, on orthologous pairs of genes (BLASTp reciprocal best hits with at least 40% identity, 70% coverage, and an E-value lower than 1e-10), were counted within a non-overlapping 100 kb window starting from *dnaA* gene (corresponding to *oriC*, approximately at position 6, 204, 383 in RLB1-8). The number of transitions in windows of the right replichore was added to the number of transitions of windows at the same distance of *dnaA* in the left replichore and the sum was normalized to the total number of coding positions of the window. A significant positive correlation was observed between the transition count and the genomic position (Kendall rank correlation test *r*=0, 38 *P*=0, 00005).

As BPS accumulation is low in the WT evolved lines, we decided to estimate the accumulation of transitions by comparing WT genomes of strains that have been diverging for much longer than 60 sporulation cycles. Therefore, the genomes of *Streptomyces* environmental strains RLB1-8 and RLB3-17 (44), isolated at a spatial microscale (on the order of one cubic millimeter) and sharing an average nucleotide identity (ANI) of 99.19% were compared in terms of SNP distribution. Orthologs were defined by pairwise comparison of both strain genomes and detection of transitions was first specifically done in orthologous pairs. As above, the number of transitions was counted within a sliding non-overlapping 100 kb window, starting from *oriC* (estimated at position 6, 204, 383 by the *dnaA* gene location in RLB1-8 genome, which was used as the reference), extending towards the left or right end of the chromosome. The sum of equidistant windows on the left and the right replichores was normalized to the total number of coding positions included in the windows (to consider the possible differential density of orthologous genes along the chromosome) and plotted as a function of the distance from *oriC.* As shown in Figure 4C, a clear increase of transition number in orthologous genes is observed when the distance from the origin of replication increases (Kendall rank correlation test *r*=0, 38 *P*=0, 00005). When considering the whole genome and not only the coding regions, the transitions are also clearly more frequent when moving away from *oriC* (Kendall rank correlation test *r*=0, 33 *P*=0, 00045) (Supplementary Figure S4). The comparison of genomes of other conspecific strains isolated from the same soil grain showed a similar increasing trend towards the chromosome termini (data not shown). Altogether, these results indicate that a short time evolutionary scale is sufficient to observe transition accumulation with a crescent-shaped distribution along the chromosome of WT strains possessing a functional non-canonical MMR.

### Occurrence of BPSs has a strong DNA-strand bias

When considering the 5’ to 3’ nucleotide sequence (“top strand”) of *S. ambofaciens* chromosome, the left replichore (starting at the left end to the *oriC* position) corresponds to the lagging strand while the right replichore (starting at *oriC* to the right end) corresponds to the leading strand. The A:T>G:C transitions occur when C is mispaired with template A or G is mispaired with template T. Considering one strand, A:T>G:C transitions are either A>G or T>C, and our results in *ΔnucS* evolved lines (Supplementary Table S4) showed that in the leading strand (right replichore), A>G substitutions were three times more frequent than T>C substitutions (841 versus 290, respectively). This bias was reversed in the lagging strand (left replichore) where mispaired Ts occurred more often than mispaired As (1153 Ts versus 702 As). This means that during replication, A:T>G:C transitions are more likely to originate from a C in the lagging strand mispaired to template A and a G in the leading strand mispaired to a template T than the contrary. Likewise, a similar strand bias was found for G:C>A:T transitions with more C>T than G>A substitutions in the right replichore (507 versus 200, respectively); the reversed trend was observed in the left replichore (442 versus 773, respectively). Thus G:C>A:T transitions are more likely to originate from a A in the lagging strand mispaired to template C and a T in the leading strand mispaired to a template G than the contrary. This transition strand bias is consistent with the observations made in MMR defective *E. coli* (4), C*. glutamicum* (21) and *M. smegmatis* (19). Since this finding was made in the mutant lines lacking a functional MMR, and since NucS-dependent MMR in *Streptomyces* was able to correct G/T but not A/C mismatches *in vitro*, the above results imply that transitions arise more frequently from a G mispaired to a T on the lagging strand template than a T on the leading strand template, which indicates a nucleotide misincorporation bias by the DNA polymerase. No strand bias could be observed in the WT evolved lines, because the mutation rate is too low. Therefore, further investigation is needed to clarify a possible strand-specific activity of the non-canonical MMR.

## DISCUSSION

*Streptomyces* species, like the majority of actinobacteria, do not possess the signature proteins of the canonical MMR pathway, MutSL. The discovery of NucS in archaea as part of an alternative MMR pathway prompted further investigation. NucS is not only found in many archaea but also widely conserved among most actinobacteria (9, 18). In our analysis, we found that NucS*_Sam_* is highly conserved within the *Streptomyces* genus, establishing *nucS* as a core gene in these bacteria. *S. ambofaciens* Δ*nucS* mutant displayed a hypermutable phenotype characterized by the appearance of sectored colonies. A sector on a colony (with a radial growth) is a visible different hyphal phenotype resulting from a mutation that occurred during the colony growth. The frequency of sectors is therefore a visual indication of mutation rate. The increased emergence of rifampicin-resistant colonies in the progeny of Δ*nucS* mutant also indicated a significantly increased spontaneous mutation rate. These results are consistent with previous studies conducted in other actinobacterial species, and provide additional evidence for the role of NucS in genome maintenance (18, 20, 21). Our biochemical study demonstrated that NucS*_Sam_* targets G/T, T/T, and G/G dsDNA substrates by cleaving both strands, resulting thus in a DSB. This cleavage mechanism and preference are in accordance with those described in archaeal species and in *C. glutamicum* (9, 20, 21). The Mn^2+^ enzymatic cofactor was required for NucS*_Sam_* activity, in the presence or absence of β-clamp*_Sam_*. However, no cleavage activity was observed with Mg^2+^ for NucS*_Sam_*. This differs from NucS from *T. kodakarensis* and *C. glutamicum* which use both Mn^2+^ and Mg^2+^ (9, 20, 21). Such activity has not yet been shown in *Mycobacterium* species, and one of the suggested hypotheses was the need for other partners (18). Indeed, the endonuclease activity of NucS*_Sam_* was significantly enhanced by the presence of the β-clamp, a crucial protein involved in DNA replication. This interaction with β-clamp is essential for efficient mutation avoidance and genome stability in *C. glutamicum* (20, 21). It has been hypothesized that the β-clamp could induce a conformational change that favors DNA binding or induce the catalytic step (17) but most importantly, it could tether the MMR enzyme to the replication zone where mismatches occur (21). The coupling of canonical MMR to the β-clamp has also been demonstrated as the interaction of MutS and MutL with β-clamp is necessary for a functional MMR in *E. coli* and in *B. subtilis* (12, 53, 54).

Our biochemical study also revealed an additional endonuclease activity of NucS*_Sam_* on deaminated substrates such as G/U and T/I mismatches. Interestingly, this activity was only observed in the presence of β-clamp. This suggests that NucS*_Sam_*, together with β-clamp, plays a role in the repair of DNA lesions resulting from deamination similar to the base excision repair (BER) pathway. Conversion of cytosine to uracil leads to G:C>A:T base pair mutation in the subsequent DNA replication. Uracil-DNA glycosylase (UDG) is a key enzyme that cleaves the N-glycosidic bond thereby initiating the BER pathway. It has been shown that UDG genes are highly conserved across *Streptomyces* genus and that the deletion of these genes is not lethal in *S. lividans* (55). In the light of these results, it is conceivable that, in addition to UDGs, NucS*_Sam_* may play a significant role in the BER-like pathway in *Streptomyces*. This hypothesis is supported by structural and biochemical investigations of NucS from the hyperthermophilic archaeon *T. gammatolerans*, which is thought to be involved in an alternative pathway for the repair of deaminated bases (51). The study has highlighted that the ends of cleaved dsDNA products could be utilized by homologous recombination proteins. However, the involvement of PCNA was not investigated in this study. Our results suggest that the activity of NucS*_Sam_* on deaminated bases is coupled to replication, as it requires the presence of β-clamp. Uracil or hypoxanthine incorporation into DNA can occur when the pool of pyrimidine or purine is depleted, leading to the erroneous incorporation of uracil or hypoxanthine instead of cytosine or adenine, respectively (56, 57). In this context, it can be speculated that NucS*_Sam_* may counteract the misincorporation of uracil and hypoxanthine by the DNA polymerase during replication. Another possibility is that NucS*_Sam_* plays a role in a mechanism similar to pre-replicative BER. Pre-replicative BER, mainly described in eukaryotes, is thought to target and repair base lesions on DNA template upstream the replication fork and this function has been linked to the DNA-glycosylase NEIL1 and to PCNA in human cells (58). Interestingly, *M. tuberculosis* Nei2, a BER glycosylase that removes oxidized base lesions from ssDNA and replication fork-mimicking substrates, has recently been shown to interact with β-clamp, which enhances its activity (59, 60). NucS*_Sam_*, through its interaction with β-clamp*_Sam_*, could have a similar role in targeting deaminated bases upstream the replication fork. However, it remains unclear whether and how NucS*_Sam_* targets deaminated base sites that arise spontaneously independently of replication, for example during oxidative stress.

A significant transition-biased spectrum was observed in Δ*nucS* strains through MA assays. This is in line with previous works in actinobacteria and archaea (19, 21, 61). Such transition shift is a well-established hallmark of MMR deficiency as observed in organisms bearing a canonical MMR such as *E. coli* (4). It is indeed known that transition mispairs are less efficiently proofread than transversions, making them specific targets of MMR machineries (62). It is important to point out that Δ*nucS* lines accumulate more A:T>G:C than G:C>A:T transitions whereas the opposite is observed in WT lines. Thus, the non-canonical MMR could prevent the accumulation of GCs in *Streptomyces* genomes, which already have a high GC content, as suggested for GC-rich mycobacterial genomes (19), and also with opposite observations in AT-rich *E. coli* genomes (4). A very interesting result is the correlation that can be made with our *in vivo* and *in vitro* findings. Indeed, *in vitro* activities revealed that the G/T mismatch is clearly the preferred substrate of NucS*_Sam_*, whereas no cleavage activity on A/C mispair could be detected. Such striking preference was also observed with NucS from *T. kodakarensis* (9) and *C. glutamicum* (20, 21). Furthermore, recent oligonucleotide recombination experiments in *M. smegmatis* yet to be published (63) revealed that *in vivo* NucS would only correct G/T but not A/C mismatches in this species. In this case, transition events in MMR-deficient *ΔnucS* lines are due to uncorrected G/T mismatches. For example, during replication, a G>A transition can arise from a mispaired G/T or a mispaired C/A (Figure 5, upper panel). Since NucS only targets G/T mismatches, the increase of G>A transition mutation in the absence of NucS is due to accumulation of G/Ts (Figure 5, lower panel), and this is also true for C>T, A>G and T>C transitions. Therefore, the drastic transition accumulation in *Streptomyces ΔnucS* lines reflects the frequency of G/T mismatch occurrence while the negligible frequency of uncorrected A/C mispair events can be estimated by the residual transition accumulation in genomes of WT lines possessing a functional non-canonical MMR. These results enable to discriminate which transition mispairing is the most frequent during incorporation of nucleotides by DNA polymerase. The replication complex is thereby prone to generate far more G/T than C/A mismatches. Our results, supported by MA and biochemical studies on *C. glutamicum* (20, 21), highlight a remarkable *in vivo* property of the DNA polymerase. The hypothesis that G*/*T mismatches are much more likely to occur *in vivo* has previously been proposed for canonical MMR-bearing bacteria, since MutS protein preferentially binds G/T over A/C (64). MA assays in yeast showed that G pairing with T is the most common mispair generated by DNA polymerase (65). Many studies on DNA polymerase fidelity have been performed *in vitro*, and recently a high throughput analysis based on next-generation sequencing showed general DNA polymerase preference for dGTP and dTTP misincorporation at T and G bases (66). This preference can be explained by tautomer occurrence and stability, and spatial constraints in the polymerase active site during catalysis of the phosphodiester bond (67–71). The DNA bases exist as tautomers (interchangeable pairs of isomers), and A/T and G/C base pairs contain each base in its predominant tautomeric form. Watson and Crick (72) proposed that if nucleotide bases adopted their energetically less favorable tautomeric forms, mismatches could arise in a Watson–Crick (WC)-like geometry and consequently lead to spontaneous mutations. However, it is only in the last decade that this 70-year-old model has been experimentally validated, and, in particular, G/T tautomers were demonstrated to exist during DNA polymerization and to have a role in base pair mis-incorporation and transition occurrence (73). Altogether, these data underline the propensity of DNA polymerases to produce G/T mispairs and our *in vivo* results together with observations in *C. glutamicum* (20, 21) demonstrate that these G/T mismatches are specifically eliminated by a NucS-dependent MMR mechanism, which has evolved to correct these frequent transition mismatches in cells.

**Figure 5.**
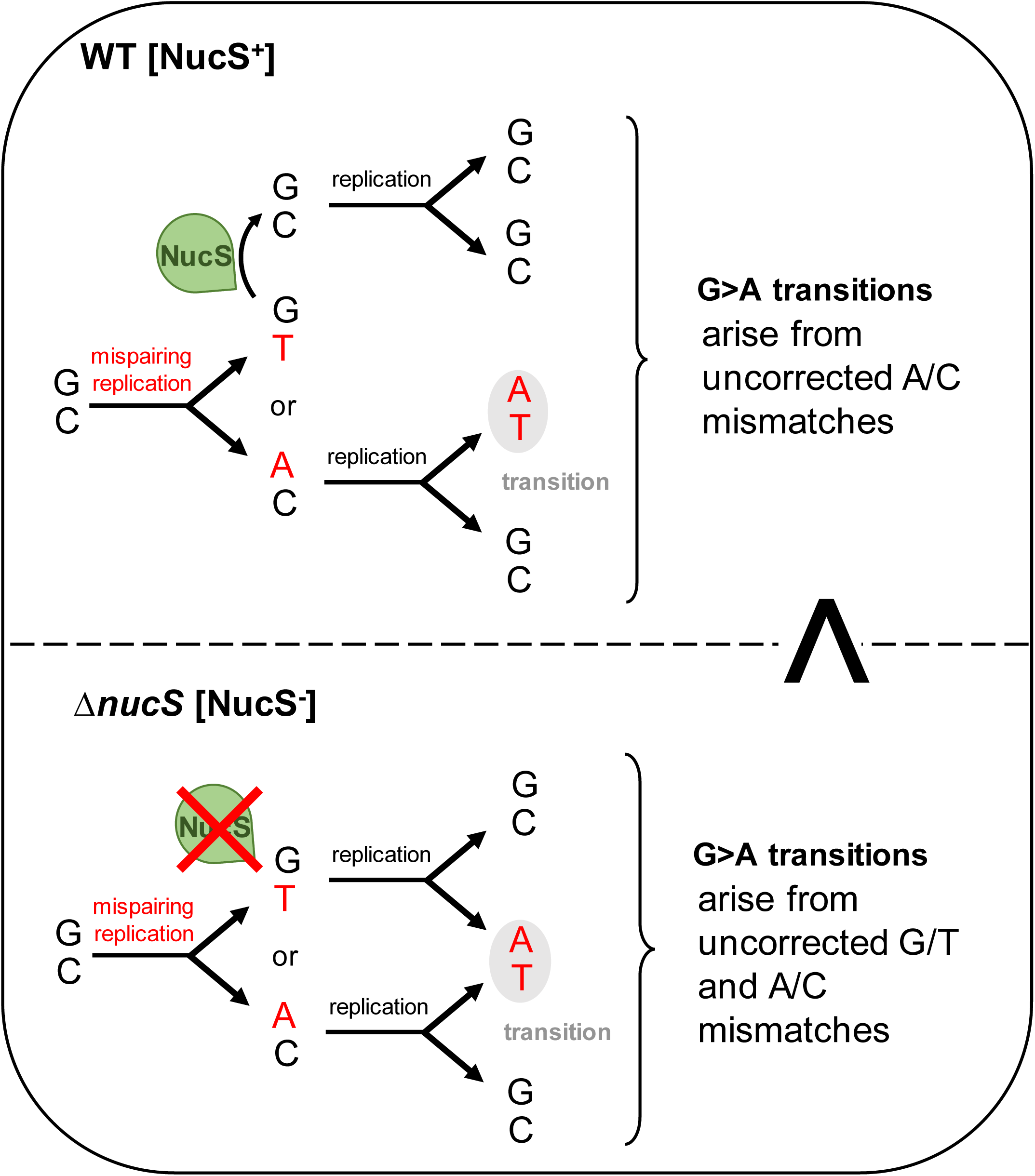
Implication of NucS mismatch preference in transition events during replication. The diagram illustrates the events leading to the base substitution of a G into an A, which is one of the four transitions that can occur on one strand during replication (the others being A>G, C>T and T>C). The sequence of events is described in WT bacteria possessing a functional non-canonical MMR (NucS endonuclease is active) and in a Δ*nucS* strain (lacking NucS-dependent MMR). Mutated Δ*nucS* lines accumulated far more G>A transitions than WT lines (shown with a superscript >).

Another remarkable observation is that accumulation of mutations in evolved *ΔnucS* lines correlates with the distance to the origin of replication. It is noteworthy that the frequency of transitions, which represent the vast majority of mutations in the mutant, follows the same trend. This crescent-shaped BPS distribution in the absence of a functional MMR reflects a variable mutation rate of DNA polymerase (due to variable accuracy or an imbalance in the nucleotide pools) and/or a differential selection along the chromosome. It is known that core genes are located in the central compartment of the chromosome in *Streptomyces* (25) and, although there is no data on the distribution of essential genes, it can be argued that most essential genes are among the core genes, so it is likely that some mutations in the central region of the chromosome are counter-selected. A commonly used way of assessing selection bias is the ratio of non-synonymous to synonymous mutations, assuming that synonymous mutations are relatively neutral. Our results showed that there was no correlation between the dN/dS ratio and the genomic position in the evolved *ΔnucS* lines. Therefore, the BPS distribution in the absence of a functional MMR reflects the distribution of mismatches that occur during replication and suggests an increasing mutation rate of the DNA polymerase towards the extremities (Figure 6). Given the propensity of DNA polymerase to generate G/T mismatches (as discussed above), this implies that G/T mispairs accumulate with the distance from the replication origin, a finding that, to our knowledge, has never been documented for bacterial linear chromosomes. Furthermore, our results of differential mutation rate along the chromosome suggest that different areas of the genome evolve at different rates in *Streptomyces*, implying that the genes located closer to the replication origin are less prone to evolution than those closer to the termini, which is consistent with the compartmentalization of *Streptomyces* chromosome (25). In WT evolved lines, the low total number of mutations prevented us from building a robust statistical model for transition distribution. However, the comparison of genomes of two close *Streptomyces* individuals isolated from a soil grain and deriving from a recent common ancestor revealed that the transition rate increases significantly with the distance from the origin. This finding suggests that this short time evolutionary scale is long enough to observe that transitions in WT environmental strains tend to accumulate in the extremities of the chromosome, where transition mismatches are more likely to occur and/or escape proofreading and MMR activities. Pioneering studies on *E. coli* and *Salmonella typhimurium* genomes indicated that genes located further from the replication origin have higher mutation rates than those closer to the origin (74, 75), and some studies suggest that it is the proximity to the fragile replication termination sites that is responsible for this phenomena (76, 77). In the last decade, MA studies on lines lacking MMR in *E. coli* or *Bacillus subtilis* (78, 79) and in *Vibrio* and *Pseudomonas* species(76) revealed a more precise wave-like pattern that is symmetrical about the replication origin. Such wave-like pattern of mutation distribution was not visible in the genome of evolved *ΔnucS* lines, either because the total number of mutations in this study is not high enough to build such a model, or because the *Streptomyces* linear chromosome has a different tridimensional structure (see below). In *E. coli*, it has been observed that intrinsic errors by the DNA polymerase occur more frequently near the replication origin, but also that error correction by the proofreading activity of the DNA polymerase or canonical MMR is more effective close to the *oriC* region. More importantly, it has been hypothesized that DNA replication becomes inaccurate in regions of high superhelical density and that nucleoid-associated protein (NAP) binding increases mutability by impeding DNA repair and/or interfering with DNA polymerase processivity during replication (77, 79, 80). *Streptomyces* NAPs are numerous and their binding along the chromosome is temporally and spatially variable throughout the bacterial life cycle (81–83) such that the association between NAPs and mutation density is complicated. An interesting example is the HU family-protein HupS which participates to chromosomal spatial rearrangement and compaction at the onset of sporulation in *Streptomyces venezuelae* (82). Increased binding of HupS to the subterminal regions organize them into a closed structure, that might be prone to a higher mutation rate.

**Figure 6.**
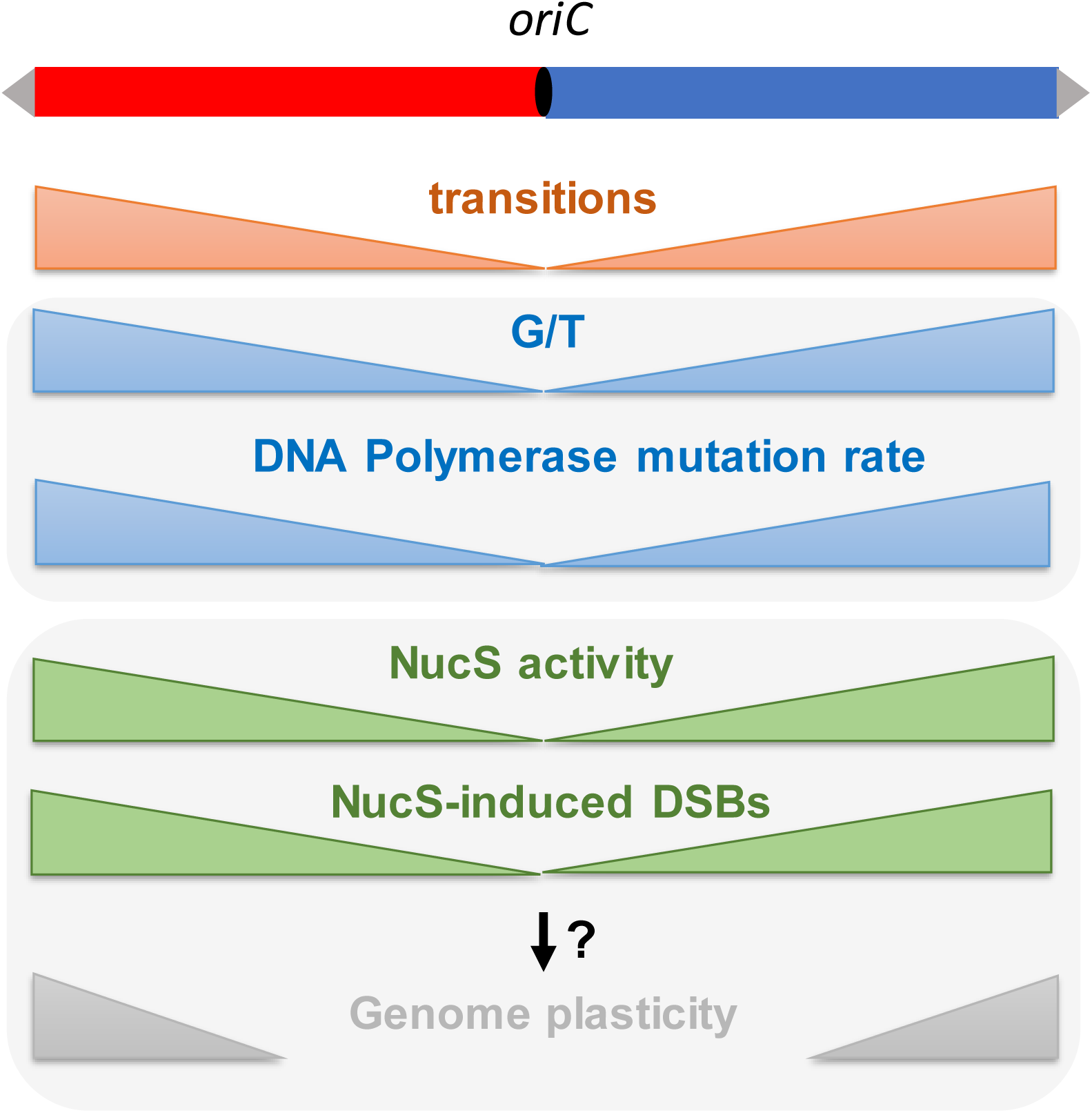
DNA polymerase mutation rate and non-canonical MMR activity along the chromosome during replication in *Streptomyces*. Red and blue boxes represent left and right replichores of the chromosome. Grey arrows represent the terminal inverted repeats at each chromosome extremity. In MMR defective (Δ*nucS*) lines, the transition density increases towards the extremities of the chromosome. Because NucS*Sam* cleaves G/T but not A/C mismatches *in vitro*, this transition trend suggests an increase of G/T mispairing and an increase in DNA polymerase mutation rate towards the extremities (in blue). This observed mutation profile also suggests that in MMR proficient (WT) lines, NucS activity increases with the distance of the replication origin, and thus NucS-induced DSBs are more frequent towards the extremities of the chromosome (in green). Assuming that these DSBs are repaired by recombination, this profile is in accordance with the recombination gradient driving the genome plasticity described in *Streptomyces* (illustrated in grey).

The mechanism by which NucS-mediated DSBs are repaired remains elusive. The involvement of homologous recombination to process the unwanted DSB is the most obvious hypothesis and would allow a mismatch repair without strand discrimination. The fact that many bacteria and archaea possessing NucS are polyploid, such as *C. glutamicum* or *T. kodakarensis* (84, 85), favors this postulate. *Streptomyces* are filamentous bacteria that share the same characteristics since they display polyploid hyphae during vegetative growth.

Furthermore, pull-down assays in *P. abyssi* have shown that NucS might interact with Hef and PCNA clamp (8). Hef proteins are known to play a crucial role in homologous recombination by resolving branched DNA structures such as Holliday junctions. In addition to NucS, PCNA clamp is also known to bind to the Mre11/Rad50 complex in *P. abyssi*, (86). The Mre11/Rad50 complex is involved in DSB repair and is known to engage the homologous recombination pathway by tethering and resecting DNA ends. This suggests that PCNA, which interacts with NucS and the Mre11/Rad50 complexes, may act as a master regulator of the MMR and homologous recombination pathways. Interestingly, in human cells, it has been proposed by Chen *et al.* (87) that PCNA loading on DSBs appearing independently of replication, and functions as a processivity factor for the Exo1 nuclease in DNA resection, thus directly linking DSB repair and homologous recombination mechanism to the sliding clamp. More recently, it was shown in *E. coli* that RecF targeting to post-replicative gaps involves a direct interaction with DnaN/β-clamp (88). Single-strand DNA post-replication gaps are frequently formed during replication and are predominantly repaired by homologous recombination. The recruitment of homologous recombination to DNA damage *via* the replication clamp is therefore a plausible hypothesis that could apply to the resolution of NucS-induced post-replicative DSBs. It should be noted that in *Streptomyces*, as in all bacteria, *dnaN* and *recF* genes are located at the same locus, close to *dnaA*, suggesting their essential role in replication. Besides, unlike most bacteria, *Streptomyces* species have the ability to repair DSBs by both homologous recombination and illegitimate recombination pathways. Our previous work has demonstrated the existence of a functional NHEJ pathway in *Streptomyces* (33). Hence, NucS-dependent DSBs may also be repaired by NHEJ in *Streptomyces*. Very recently, a preprint manuscript (63) suggested that mycobacterial NucS participates in a MMR reaction involving short patch repair confined within 5-6 bp of the mispaired nucleotides, which appears to be mechanistically distinct from repair by homologous recombination or NHEJ DSB repair mechanisms. A similar study in *M. smegmatis*, not approved by peer review yet, has also shown that a NucS-promoted DSB is processed by a 5’- 3’ exonuclease, but is independent of both RecA, RadA, and NHEJ functions Ku and LigD (89).

The high level of genetic instability observed in the *Streptomyces* genus is one of its most intriguing features and DSB repair is thought to be the main driver of this instability (31). The increasing occurrence of mismatches towards the ends of the chromosome suggests that NucS activity is more required further from the origin and consequently that more DSBs occur in the subtelomeric regions (Figure 6). Several studies have pointed to the existence of a recombination gradient towards the chromosomal arms highlighting the compartmentalization of the *Streptomyces* genome (25, 44). Our results support an increase of DSBs towards the ends of the chromosome, contributing to increased recombination events in the extremities and thus to the unique organization of the *Streptomyces* chromosome. Yet, how the NucS-dependent DSBs are repaired remains to be elucidated. Additional aspects such as differential efficiency of DSB repair along the chromosome as well as the involvement of the spatial conformation of the chromosome (90) or the role of chromatin-organizing proteins (82) need to be considered to gain a comprehensive understanding of *Streptomyces* genome dynamics.

## DATA AVAILABILITY

The sequencing data underlying this article have been deposited with links to BioProject accession number PRJNA1047308 in the NCBI BioProject database (https://www.ncbi.nlm.nih.gov/bioproject/).

## ACKNOWLEDGEMENTS

The authors would like to acknowledge the ASIA platform and Tiphaine Dhalleine for the technical expertise with SEC-MALS analyses.

