## Supplementary data for "Correction of non-random mutational biases along a linear bacterial chromosome by the mismatch repair endonuclease NucS"

**Supplementary Table S1. Primers used in this study.**

| Name | Sequence (5'-3') | Purpose |
|---|---|---|
| nucS_compl_F | AGGGGAATTCAGACGTCGAGCGACATGGGT | Cloning *nucS* in pSET152 for the creation of complemented strains |
| nucS_compl_R | CGAGGAATTCGACGGAGCAGATCATCTGAGC | |
| nucS_F | GTTGCCGCATATGCGTCTCGTCATTGCCCGCT | Cloning *nucS* in pET15b for heterologous expression |
| nucS_R | TGTAGGATCCTCAGAACAGCCGCAGCTTG | |
| dnaN_F | TAGCATATGAAGATCCGGGTGGAACGC | Cloning *dnaN* in pET15b for heterologous expression |
| dnaN_R | ACTGGATCCTCAGCCGCTCAGCCGCAC | |

**Supplementary Table S2. Sequence of oligonucleotides used for cleavage assays (left) and combination of the oligonucleotides to form heteroduplex DNA with a mismatch (right).**
Asterisk means that the sequence is 6FAM labeled at 5′

| Number | Sequence (5'-3') |
|---|---|
| 1* | ACCCAGTACTCGCTGCTGAATGGAGCCGCGCGGCTGAAGGACA |
| 2* | TGTCCTTCAGCCGCGCGGCTCCGTTCAGCAGCGAGTACTGGGT |
| 3 | TGTCCTTCAGCCGCGCGGCTCCATTCAGCAGCGAGTACTGGGT |
| 4 | TGTCCTTCAGCCGCGCGGCTCCGTTCAGCAGCGAGTACTGGGT |
| 5 | TGTCCTTCAGCCGCGCGGCTCCTTTCAGCAGCGAGTACTGGGT |
| 6 | TGTCCTTCAGCCGCGCGGCTCCCTTCAGCAGCGAGTACTGGGT |
| 7 | TGTCCTTCAGCCGCGCGGCTCCAATCAGCAGCGAGTACTGGGT |
| 8 | TGTCCTTCAGCCGCGCGGCTCCACTCAGCAGCGAGTACTGGGT |
| 9 | ACCCAGTACTCGCTGCTGAAGGGAGCCGCGCGGCTGAAGGACA |
| 10 | ACCCAGTACTCGCTGCTGAAAGGAGCCGCGCGGCTGAAGGACA |
| 11 | ACCCAGTACTCGCTGCTGAACCGAGCCGCGCGGCTGAAGGACA |
| 12 | ACCCAGTACTCGCTGCTGAAUGGAGCCGCGCGGCTGAAGGACA |
| 13 | TGTCCTTCAGCCGCGCGGCTCCITTCAGCAGCGAGTACTGGGT |

| Mismatch | Combination |
|---|---|
| none | 1+3 |
| G/T | 1+4 |
| T/T | 1+5 |
| T/C | 1+6 |
| A/A | 1+7 |
| A/C | 1+8 |
| G/G | 2+9 |
| G/A | 2+10 |
| C/C | 2+11 |
| G/U | 2+12 |
| T/I | 1+13 |

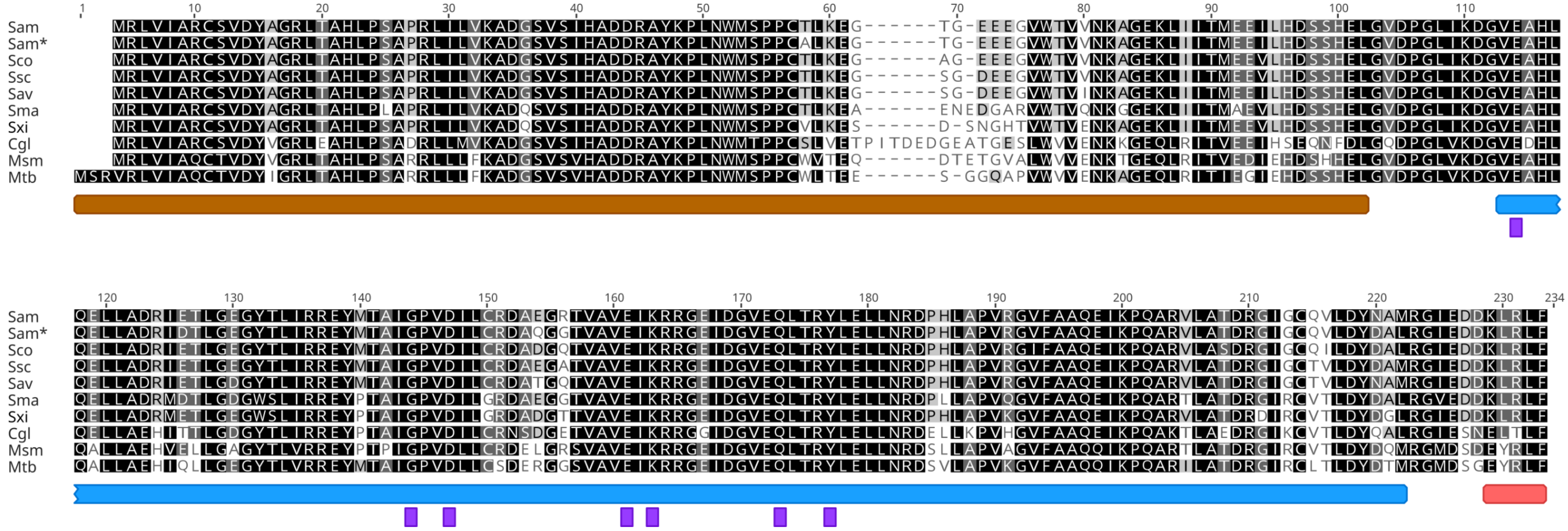


**Supplementary Figure S1. Conservation of NucS protein through actinobacteria.** Alignment of NucS protein sequences from different species of *Streptomyces* and *Mycobacterium*. *S. ambofaciens* ATCC 28777 (Sam), *S. ambofaciens* DSM 40697 (Sam*), *S. coelicolor* A3(2) (Sco), *S. scabiei* NCPPB 4086 (Ssc), *S. avermitilis* MA 4680 (Sav), *S. xiamenensis* 318 (Sxi), *S. marincola* strain SCSIO 03032 (Sma), *C. glutamicum* ATCC 13032 (Cgl), *M. smegmatis* $mc^2$ 155 (Msm), *M. tuberculosis* H38rv (Mtb). Black, dark grey, light grey and blank boxed residues correspond to 100%, 80-100%, 60-80% and less than 60% sequences presenting the same residue. Colored boxes under the alignment indicate protein domains according to *Pyrococcus abyssi* NucS structure (Ren et al., 2009). DNA-binding and catalytic domains are shown in brown and blue respectively. PIP-box motif is represented in red. Essential residues for nuclease activity are shown in purple.

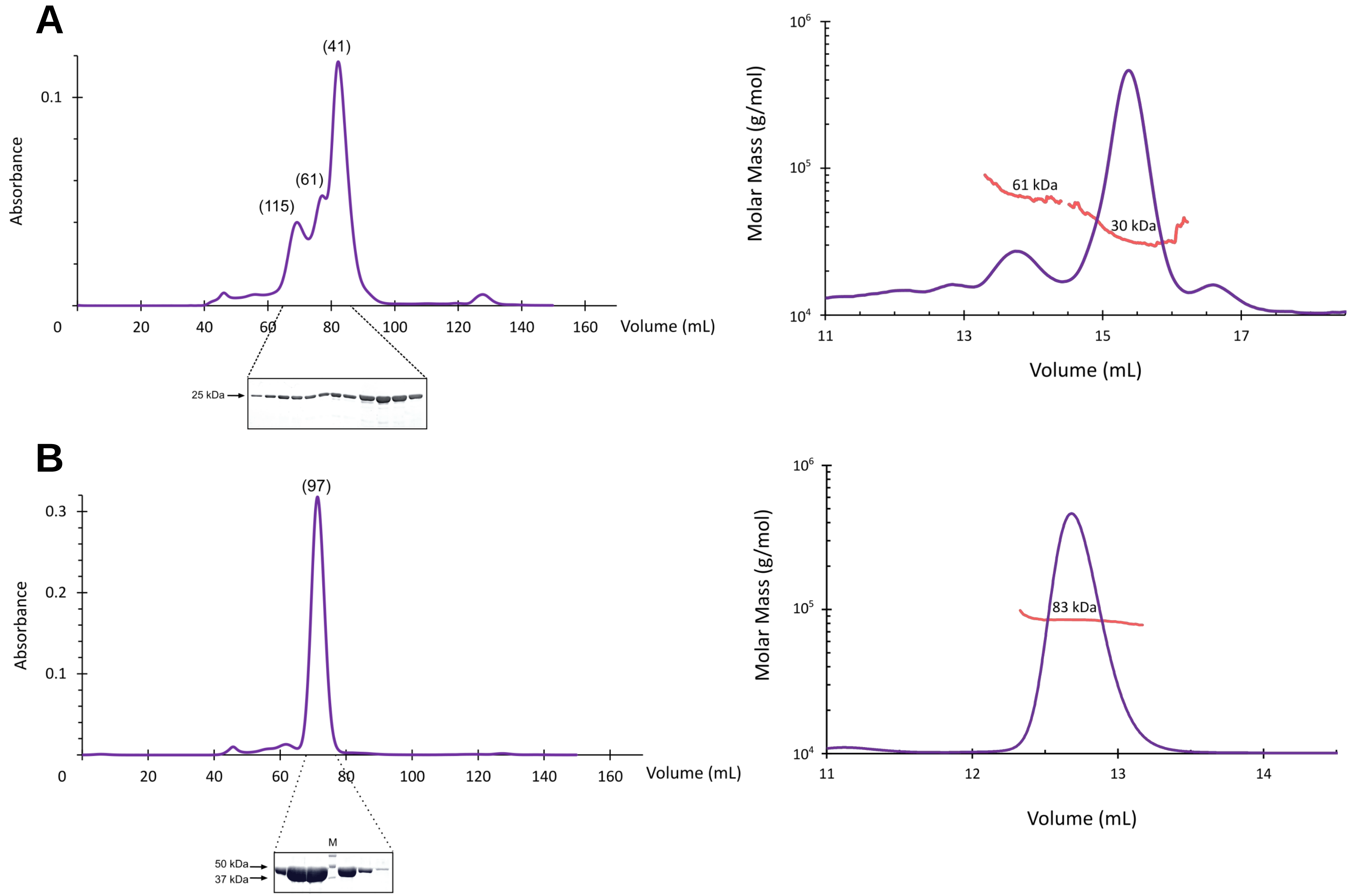


**Supplementary Figure S2. Purification and oligomeric states of NucS$_{Sam}$ and β-clamp$_{Sam}$** Gel filtration profile and SEC-MALS analysis (left and right panels, respectively) of (A) NucS$_{Sam}$ and (B) β-clamp$_{Sam}$. For gel filtration, the samples were loaded onto the Superdex 200 16/600 column. Number in brackets correspond to estimated molecular weight (kDa) using calibration curve. Aliquots of the 12 elution fractions from 63 to 87 mL for NucS$_{Sam}$ and the 6 elution fractions from 67 to 79 mL for β-clamp$_{Sam}$ were subjected to 12% SDS-PAGE (lower part). M is the protein ladder "Precision Plus Protein Standard" (Biorad). The SEC-MALS chromatograms display the UV absorbance at 280 nm and red lines indicate the molar mass distribution. Because of the reasonable homogeneity of NucS$_{Sam}$ peaks, the molecular weight was estimated at the right extremity of each peak.

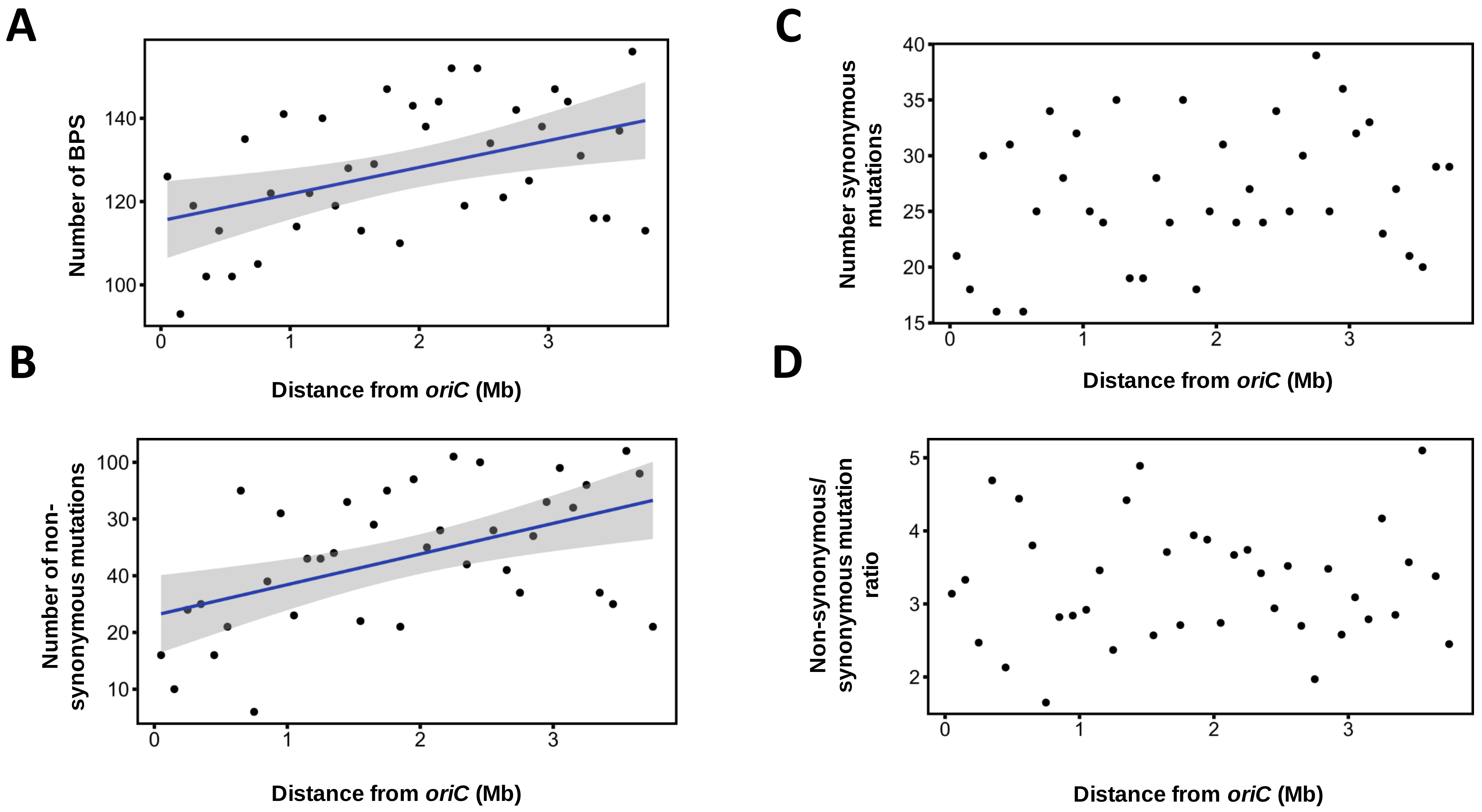


**Supplementary Figure S3. Distribution of mutations along the chromosome in Δ*nucS* lines.** Base pair substitutions (BPSs), non-synonymous and synonymous mutations within coding sequences, were counted within a non-overlapping 100 kb window, starting from *dnaA* gene (located in the middle of the chromosome at position 4,021,374 and approximately corresponding to the origin of replication, *oriC*) and sliding towards the chromosome extremities. The number of these mutations in windows of the right replichore were added to the number in windows at the same distance of *dnaA* in the left replichore. The sum of BPSs (A), non-synonymous mutations (B), or synonymous mutations (C) is represented as a function of the distance from *dnaA*. The ratio of non synonymous to synonymous mutations (dN/dS) was calculated for each 100 kb window and plotted as a function of the genomic position (D). Significant positive correlations were observed between the genomic position and the BPS count (Pearson's correlation coefficient $r$=0.451, $P$=0.004), or between the genomic position and the non-synonymous mutations (Pearson's correlation coefficient $r$=0.499, $P$=0.001). No correlation was noted for synonymous mutations or for dN/dS ratio.

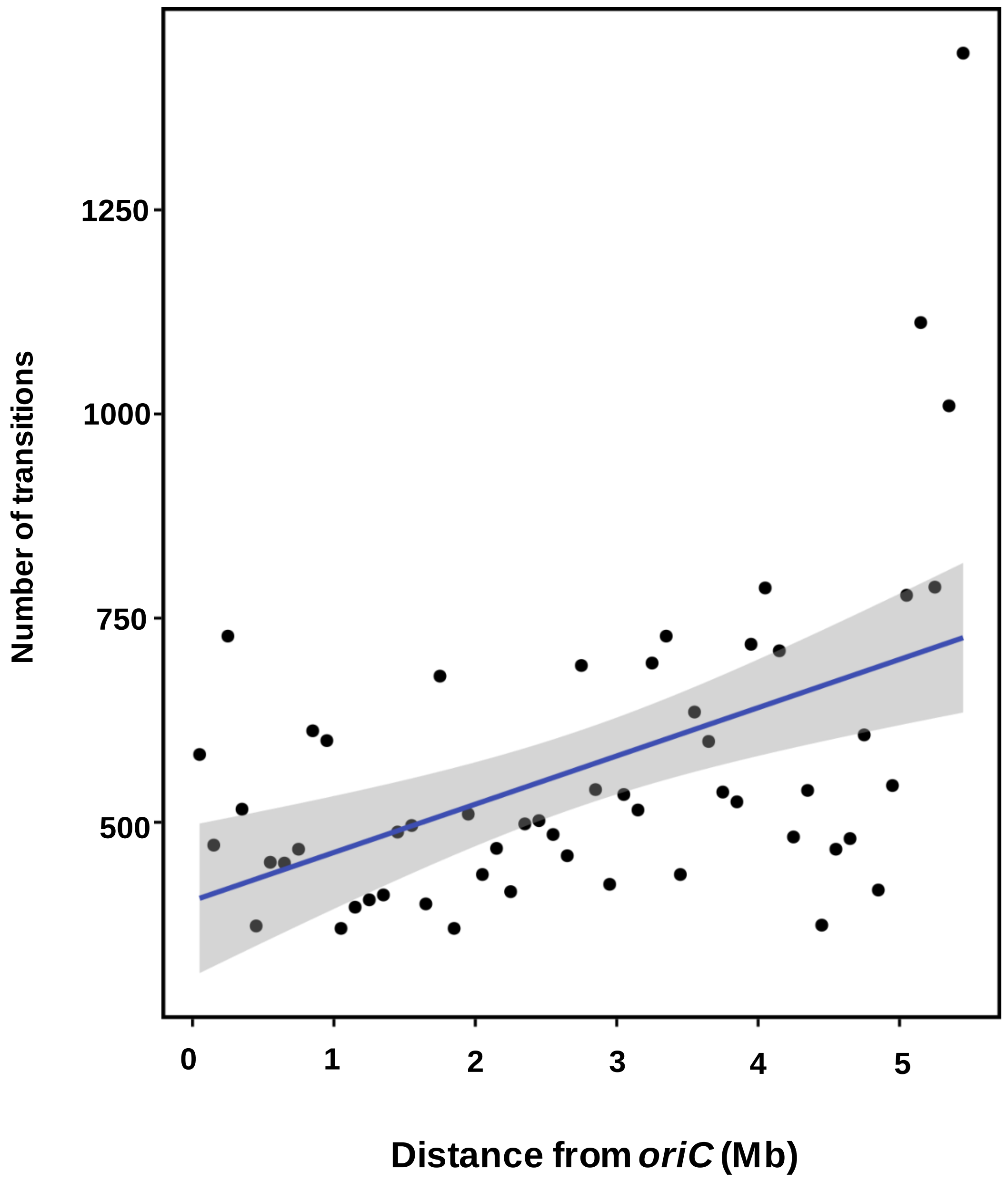


**Supplementary Figure S4. Transition distribution along the chromosome of *Streptomyces* environmental strain RLB1-8 compared to *Streptomyces* environmental strain RLB3-17.** Transitions were counted within a non-overlapping 100 kb window starting from *dnaA* gene (corresponding to *oriC*, approximately at position 6,204,383 in RLB1-8). The number of transitions in windows of the right replichore was added to the number of transitions of windows at the same distance of *dnaA* in the left replichore , and represented as a function of the distance from *dnaA* gene. A significant positive correlation was observed between the transition count and the genomic position (Kendall rank correlation test *r*=0,33 *P*=0,00045).

**Supplementary Table S4. Distribution of BPSs in the replichores**

| | WT | | Δ*nucS* | |
|---|---|---|---|---|
| | Left Replichore | Right Replichore | Left Replichore | Right Replichore |
| **Transitions** | **30** | **43** | **2357** | **2551** |
| A:T>G:C | 11 | 21 | 1424 | 1562 |
| A -> G | 3 | 14 | 393 | 1150 |
| T -> C | 8 | 7 | 1031 | 412 |
| G:C>A:T | 19 | 22 | 933 | 989 |
| G -> A | 16 | 12 | 699 | 274 |
| C -> T | 3 | 10 | 234 | 715 |
| **Transversions** | **36** | **38** | **47** | **93** |
| A:T>T:A | 1 | 4 | 1 | 11 |
| A -> T | 1 | 2 | 0 | 7 |
| T -> A | 0 | 2 | 1 | 4 |
| A:T>C:G | 2 | 4 | 8 | 12 |
| A -> C | 1 | 1 | 7 | 2 |
| T -> G | 1 | 3 | 1 | 10 |
| G:C>T:A | 12 | 6 | 14 | 18 |
| G -> T | 11 | 2 | 10 | 5 |
| C -> A | 1 | 4 | 4 | 13 |
| G:C>C:G | 21 | 24 | 24 | 52 |
| G -> C | 11 | 2 | 9 | 29 |
| C -> G | 10 | 22 | 15 | 23 |
| **Total** | **66** | **81** | **2404** | **2644** |
